# Replication stress underlies genomic instability at CTCF/cohesin-binding sites in cancer

**DOI:** 10.1101/2023.10.24.563697

**Authors:** Elangoli Ebrahimkutty Faseela, Dimple Notani, Radhakrishnan Sabarinathan

## Abstract

CCCTC-binding factor (CTCF) and cohesin play a significant role in the formation of chromatin loops and topologically associating domains (TADs), which influence gene expression and DNA replication. CTCF/cohesin-binding sites (CBSs) present at the loop anchors and TAD boundaries are frequently mutated in cancer; however, the molecular mechanisms underlying this remain unclear. Here, we investigate whether the binding of CTCF/cohesin on DNA imposes constraints on DNA replication, leading to replication stress and genomic instability. Our results reveal that CTCF and cohesin remain co-bound to DNA during replication (S phase) in cancer cells (HeLa). Further, examination of replication stress through ChIP-seq of the DNA damage response/repair proteins (MRE11, STN1, γH2AX, and RAD51) showed high enrichment of these proteins at CBSs (as compared to their immediate flanking regions and control sites) and positively correlated with the binding strength of CTCF/cohesin at CBSs in the S phase. Moreover, analysis of somatic mutations from cancer genomes supports that the enrichment of mutations at CBSs is significantly higher in samples harbouring somatic copy number deletion in MRE11 and STN1 compared to wild-type samples. Together, these results demonstrate that the co-binding of CTCF/cohesin on the DNA during the S phase causes replication stress and DNA strand breaks, and this could lead to genome instability at CBSs observed in cancer.

## Introduction

Chromatin regulates the accessibility of DNA to various biological processes, including DNA replication, DNA damage and repair, and gene expression[1–3]. In eukaryotes, chromatin is organised into distinct structural units called topologically associating domains (TADs)[4]. TADs are composed of genes and their cis-regulatory elements that show frequent physical interactions between them through chromatin looping. This looping is mediated by two main architectural proteins: CTCF (CCCTC-binding factor) – a zinc finger protein that binds to its highly conserved motif sequences on the DNA; and cohesin – a ring-shaped protein complex that extrudes the chromatin loop until it encounters convergent CTCF bound motifs[5,6]. Thus, CTCF binding and its motif orientation on the DNA determine the positioning of cohesin to create loop anchors and TAD boundaries[7]. Moreover, the binding of CTCF at TAD boundaries can act as an insulator, limiting the chromatin interactions between genes and their regulatory regions predominantly within the TAD, thereby controlling expression of genes within the TAD[8,9].

Previous studies have shown that the CTCF/cohesin-binding sites (CBSs; i.e., DNA regions bound by both CTCF and cohesin), enriched at the boundaries of chromatin-loops and TADs, exhibit an increased somatic mutation rate in multiple tumour types[10–13]. However, the molecular mechanism underlying this observation is not well understood. Since the somatic mutations observed at CBSs are predominantly generated by the mutational processes active in the respective tumour type[12,13], we hypothesise that this enhanced activity of mutational processes at the CBSs could be influenced by the interplay between endogenous process (DNA replication) and binding of CTCF/cohesin to DNA. Recent studies have shown that the boundaries of chromatin-loops and TADs experience topological stress and undergo DNA double-strand breaks due to the activity of Topoisomerase II beta (TOP2B)[14,15]. Moreover, increased cohesin loading on chromatin affects replication fork progression[16] and causes replication stress in a CTCF-dependent manner in oncogenic c-MYC induced conditions[17]. Similarly, in yeast, cohesin has been shown to induce topological stress-dependent DNA damage during replication[18]. While these findings point to the topological stress imposed by the binding of CTCF and/or cohesin on the DNA, their association with replication stress and genomic instability locally at CBSs remains uncharacterized.

In this study, we found that CTCF and cohesin remain co-bound to the DNA during the replication (S) phase and are associated with replication stress in cancer cells (HeLa). Further, we show that the DNA damage response (DDR) proteins, MRE11, STN1 and FANCD2, associated with fork-stalling are enriched at CBSs during the S phase, with and without external replication stress. Moreover, we observed an enrichment of γH2AX, RAD51 and ATM at the CBSs, indicating that these sites undergo DNA double-strand breaks and repair. Finally, by using somatic mutation data from whole-genome sequencing of patient tumours (PCAWG[19] cohort), we demonstrate an elevated mutation rate at CBSs, particularly in tumours harbouring somatic copy number deletions in DDR genes (such as MRE11 and STN1) as compared to tumours without deletions in those genes. Taken together, our results reveal that the binding of CTCF/cohesin proteins to the DNA during the replication phase imposes constraints on the progression of the DNA replication fork, leading to fork-stalling and DNA strand breaks. The subsequent activation of error-prone replication or repair could lead to increased somatic mutations at CBSs.

## Results

### CTCF/cohesin co-bound to DNA during the replication phase

Although the dynamics of cohesin loading onto the chromatin has been studied during various stages of the cell cycle[20–22], the binding of CTCF to DNA and its association with cohesin during DNA replication or synthesis (S) phase is less understood. To determine whether the CTCF and cohesin proteins are bound to the DNA during S phase, we used cervical cancer cell line (HeLa) to measure the expression level of the above proteins and their genome-wide binding. Initially, HeLa cells were synchronised to get cells at different stages of cell cycle (G1/S, Early S, Mid S, Late S, Mitotic) by using thymidine arrest/release or nocodazole block method (see Methods). The extent of synchronisation under each stage was verified by assessing the cell cycle profiles using flow cytometry with Propidium Iodide (PI) staining (Fig 1A, see Methods). In addition, S phase synchronisation of the cells was verified through immunostaining for the replication marker, proliferating cell nuclear antigen (PCNA) [23,24], which exhibited a distinct staining pattern at various S phase stages (Fig S1A). The PCNA staining showed a punctate signal in 86% of the Mid S phase synchronised cells, as compared to 21% in asynchronous cells (in which the majority of the cells are in the G1 phase; Fig S1B). We also assessed cyclin expression levels in each synchronised cell population. As expected, Cyclin A showed high expression throughout the S phase, whereas Cyclin B was predominantly present in mitotic cells (Fig S1C).

**Fig 1.**
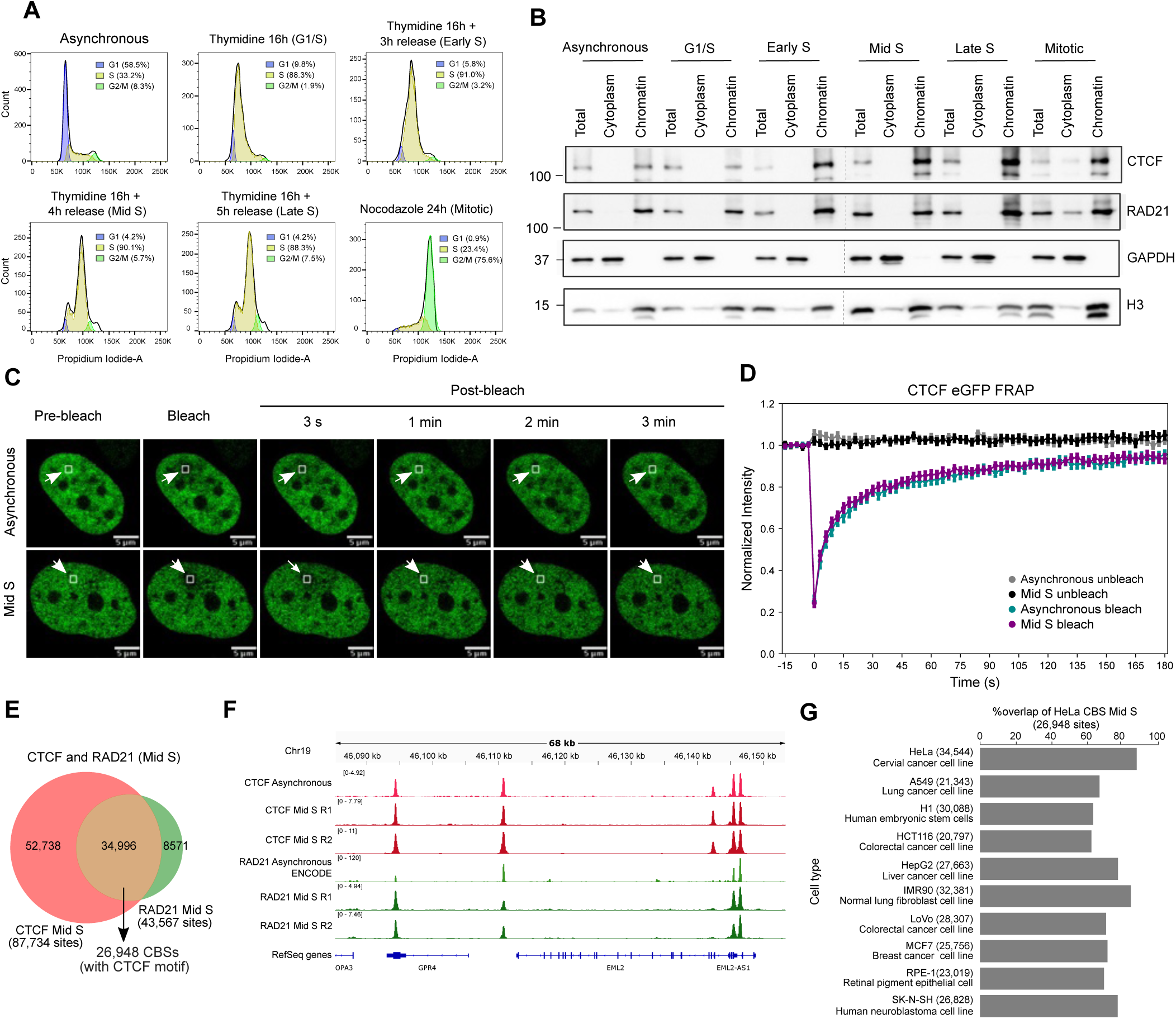
CTCF and cohesin co-bound to DNA in the replication phase. (**A**). HeLa cells were synchronised to various phases of the cell cycle by using thymidine arrest/ release or nocodazole block method (see Methods). The cell cycle profile in each stage was confirmed by staining with Propidium Iodide (PI) and analysed on a BD FACSVerse flow cytometer. In each histogram, cell count was plotted against DNA content measured from the PI fluorescence signal. The percentage of cells in each phase of the cell cycle is mentioned in brackets. (**B**) The chromatin-bound fraction of CTCF and RAD21 at various cell cycle phases of HeLa cells was checked by western blotting. GAPDH and Histone H3 were used as a loading control for cytoplasmic and chromatin-bound fractions, respectively. (**C**) Representative FRAP images of GFP-tagged CTCF in asynchronous (top panel) and Mid S (bottom panel) HeLa cells. The scale bar is 5μm. **(D)** Fluorescence recovery curve of GFP-tagged CTCF for 3 minutes post-bleaching in asynchronous and Mid S HeLa cells (37 and 41 cells in asynchronous and S phase, respectively) from 3 biological replicates. The normalised intensity value is plotted on the y-axis (each dot in the graph represents mean±1SD). (**E)** Venn diagram showing the overlapping peaks of CTCF and RAD21 in Mid S HeLa cells, and the number of high-confident CTCF/cohesin-binding sites (CBSs) with presence of CTCF motif (predicted within the CTCF peak) and RAD21 peak within 100bp from the CTCF motif (see Methods). **(F)** An example genome browser snapshot of a region on chr19 showing the overlap between ChIP-seq peaks in asynchronous and Mid S cells. From top to bottom: ChIP-seq signal in CTCF asynchronous, CTCF Mid S (two replicates), RAD21 asynchronous (from ENCODE) and RAD21 (two replicates) in Mid S HeLa cells. **(G)** Overlap of CBSs in the S phase HeLa with CBSs defined from various cell lines in asynchronous condition (from ENCODE). The number of CBSs identified in each cell line is mentioned in brackets.

Further, we performed chromatin fractionation followed by western blotting to check the endogenous level of chromatin-bound CTCF and RAD21 (a cohesin subunit) at different stages of cell cycle (Fig 1B). This showed that both CTCF and RAD21 were enriched in the chromatin-bound fraction (insoluble) at various time points of the replication phase (Early S, Mid S and Late S), similar to asynchronous cells, suggesting that CTCF and cohesin are consistently bound to chromatin across cell cycle phases, including S phase. Further, to check if binding dynamics of CTCF in S phase is different from asynchronous cells[25], we performed Fluorescence Recovery After Photobleaching (FRAP) on GFP-tagged CTCF (CTCF-eGFP) (Fig 1C, see Methods). We observed that the fluorescence recovery rate of CTCF in the S phase (median half-time recovery t_1/2_= 3.45 seconds, IQR = 1.8 seconds) was similar to that in the asynchronous cell population (median t_1/2_ = 3.46 seconds, IQR = 1.7 seconds) (Fig 1D and Fig S1D), suggesting that CTCF dynamics does not change in S phase.

Next, to determine which genomic regions are bound by CTCF/cohesin during replication, we performed chromatin immunoprecipitation followed by sequencing (ChIP-seq) for CTCF and RAD21 in the Mid S phase HeLa cells. We identified 87,734 and 43,567 DNA binding sites (peaks) for CTCF and RAD21, respectively, in the Mid S phase (Figs S1F-S1G, see Methods). Interestingly, the majority (80%) of RAD21 peaks overlapped with CTCF peaks in the S phase (Figs 1E and 1F, Figs S1E, S1H and S1I). Moreover, around 83% of RAD21 and 62% of CTCF peaks in the S phase overlapped with RAD21 and CTCF peaks from the asynchronous (G1 enriched) cells[26] (Figs S1J and S1K), suggesting that the genome-wide binding of CTCF and cohesin is broadly consistent across interphase. Of the additional CTCF peaks seen in the S phase, around 45% were proximal (< 5 kb) to existing peaks from the asynchronous cells , whereas the remaining were distributed distally, likely due to the redistribution of CTCF binding during replication (Fig S1L).

To obtain the specific DNA regions bound by both CTCF/cohesin (henceforth, CBSs) in the S phase, we focused on CTCF peaks which have a CTCF motif (see Methods and Fig S1M) and also have a RAD21 peak within 100 bp from the CTCF motif. This resulted in 26,948 CBSs (out of the total 34,996 CTCF and RAD21 overlapping peaks) in the S phase of HeLa cells (Fig 1E). Around 88% of these CBSs in the S phase overlapped with the CBSs in asynchronous cells of HeLa (Fig 1G). Also, when we compared the CBSs from HeLa S phase with interphase CBSs of other cell lines (different tissue-of-origins, from ENCODE[26]), we observed 62% to 84% overlap, indicating that these sites are consistently bound by CTCF and cohesin across multiple tissue types. Taken together, our results suggest that the CTCF and cohesin remain co-bound to the DNA throughout the interphase, including the replication phase.

### Enrichment of replication stress markers at CTCF/cohesin-binding sites in the replication phase

Next, we asked whether the binding of CTCF/cohesin on the DNA can affect the replication fork progression, leading to fork stalling and replication stress. To check this, we examined the enrichment of DNA replication stress-associated proteins (MRE11 and STN1) at the CBSs by using co-immunostaining and ChIP-seq approaches. MRE11 is a component of the MRN (MRE11-RAD50-NBS1) complex which acts as an initial DNA damage sensor and regulates DNA repair activity in response to replication fork stalling and DNA double-strand breaks (DSBs)[27]. STN1 (a part of the human CST: CTC1-STN1-TEN1 complex) is known to protect the stalled replication fork from excessive MRE11-mediated degradation[28].

At first, we performed co-immunostaining of CTCF with MRE11 and STN1 separately in asynchronous and Mid S HeLa cells to check for their colocalization by using super-resolution microscopy imaging (LSM980 with Airyscan 2, see Methods) (Figs S2A-S2H). The colocalization was measured using two different approaches: Pearson’s Correlation Coefficient (PCC)[29] which measures the linear relationship between the signal intensities of two proteins, and Mander’s Colocalization Coefficient (MCC; tM1 for channel 1 and tM2 for channel 2)[30] which measures the co-occurrence of signals in one channel with another[31]. Both approaches revealed that the CTCF and MRE11 colocalization was significantly higher in the S phase as compared to asynchronous cells (Mann-Whitney U test, two-sided, P = 2.21x10^-07^, 1.05x10^-22^, 1.44x10^-24^ for PCC, tM1 and tM2, respectively) (Figs S2B-S2D). Similarly, CTCF and STN1 colocalization was significantly higher in the S phase as compared to asynchronous cells (PCC: P=6.99x10^-03^, tM1: P=2.35x10^-10^, tM2: P=4.31x10^-19^, Figs S2F-S2H). Moreover, the colocalization of MRE11/STN1 with RAD21 was also higher in the S phase than in asynchronous cells (Figs S2I-S2P).

Further, to test the protein overlap seen by immunostaining to chromatin level (genome-wide enrichment of MRE11 at the CBSs), we performed ChIP-seq of MRE11 in Mid S HeLa cells. We examined the MRE11 coverage signal at a 10kb (+/-5kb) window centred on the CBSs in the S phase and control sites (Fig 2A). To check if this signal is non-random, we used two different control sites for comparison: a) CTCF-unbound (that is, CTCF motif predictions without ChIP-seq signals) to match for GC content, and b) random sites, where genomic regions (without CTCF motif or ChIP-seq signals) were randomly sampled with size equal to the CBSs. Interestingly, MRE11 displayed a higher enrichment at positions near CBSs while CTCF-unbound and random regions showed a lower and evenly distributed binding (Figs 2A and S2Q). For STN1, due to the unavailability of ChIP-grade antibody, we used the publicly available ChIP-seq data of STN1 with Myc-tag in the HeLa cells during the S phase upon replication damage with Hydroxyurea (HU)[32]. Similar to MRE11, we observed high enrichment of STN1 at CBSs relative to flanking regions and control sites (Figs 2B and S2R).

**Fig 2.**
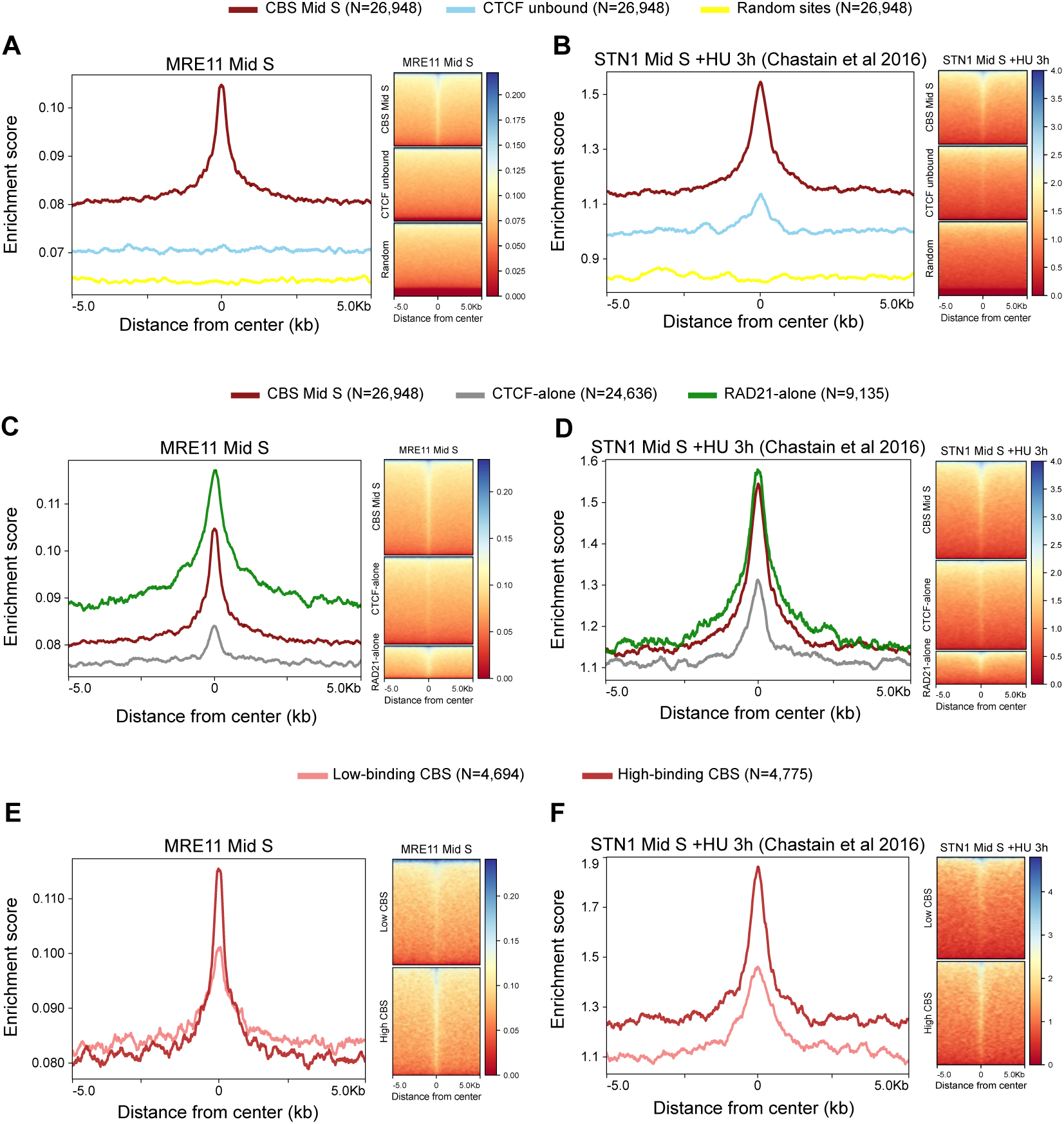
Enrichment of replication stress associated proteins at CTCF/cohesin-binding sites. **(A)** ChIP-seq profile of MRE11 in Mid S HeLa cells plotted across a +/-5kb window centred at CBSs from Mid S HeLa (maroon), CTCF unbound sites (sky blue) and random genomic regions (yellow). 26,948 sites were in each category. **(B)** Similarly, the ChIP-seq profile of STN1 in Mid S upon HU treatment (from Chastain et al. [32]) plotted surrounding CBSs and control sites (+/-5kb) **C,D:** ChIP-seq signals of **(C)** MRE11 and **(D)** STN1 (+HU) in Mid S phase plotted surrounding CBSs (maroon; 26,948 sites), CTCF-alone sites (grey; 24,636 sites) and RAD21-alone sites (green; 9,135 sites). **(E)** MRE11 profile at CBSs based on CTCF and RAD21 binding strength. Light red: CBSs with both CTCF and RAD21 low-binding (4,694 sites). Dark red: CBSs with both CTCF and RAD21 high-binding CBSs (4,775 sites). **(F)** STN1(+HU) occupancy in the Mid S phase at CTCF/RAD21 low- or high-binding CBSs spanning a +/-5kb window.

Given that the MRE11 and STN1 proteins are also involved in DNA repair, we performed the ChIP-seq analysis of another replication stress-associated protein called Fanconi anaemia D2 (FANCD2). FANCD2 is a key component of the Fanconi anaemia (FA) pathway which is crucial for interstrand crosslink (ICL) repair and also for replication stress response [33,34]. Similar to MRE11 and STN1, FANCD2 occupancy was enriched at CBSs as compared to the control sites (Fig S2S). To determine if this enrichment at CBSs is related to chromatin accessibility, we compared the ChIP-seq signals at CBSs with the DNase I hypersensitivity sites (DHSs or open chromatin regions). This showed that MRE11, STN1 and FANCD2 ChIP-seq signals were higher at the centers of CBSs compared to the DHSs (Figs S3A-S3C). However, this was not the case in the flanking regions. This suggests that the relative increase in MRE11 (STN1 and FANCD2) signals in the flanking regions of CBSs, as compared to control sites, can be attributed to chromatin accessibility around CBSs.

We then inquired whether the individual binding of CTCF or cohesin (RAD21), or their combined presence, could influence the replication stress. For this, we checked the MRE11, STN1, and FANCD2 ChIP-seq signals at CTCF-alone and RAD21-alone sites compared to CBSs (where CTCF and RAD21 co-bound). This showed a higher signal for these proteins at CBSs compared to CTCF-alone sites (Figs 2C-2D and Fig S2T), indicating that the stress at the CBSs could be associated with CTCF and cohesin co-binding. We further extended this analysis by checking the occupancy at CBSs based on the binding strength of CTCF and RAD21 together (Figs S3D and S3E). For this, we ranked the CBSs based on the binding strength of both CTCF and RAD21 at CBS and considered the top and bottom tertiles as high-binding and low-binding CBSs, respectively (see Methods). For all these proteins (MRE11,STN1 and FANCD2), we observed a relatively higher enrichment at high-binding CBSs compared to low-binding CBSs (Figs 2E-2F and Fig S2U), suggesting the binding strength of CTCF and cohesin (RAD21) is positively associated with the level of replication stress.

In addition, as compared to CBSs, RAD21-alone binding sites exhibited a higher signal for MRE11 (Fig 2C), STN1 (Fig 2D) and FANCD2 (Fig S2T). Majority of these RAD21-alone sites are located in distal intergenic regions (42%) and intronic regions (27%), and they are located far (>10 Kb) from CTCF binding sites (see Figs S3F-S3H). We then asked whether the DNA sequence composition at these sites are likely to affect replication fork progression, such as non-B DNA-forming sequence motifs or chromosomal fragile sites (CFS) (see Methods). This showed that indeed a substantial portion of RAD21-alone sites (88%) overlapped with non-B DNA-forming sequence motifs or CFS, in contrast to CBS (42%) or control sites (CTCF-unbond - 41% and random sites - 39%) (Fig S3I). Further analysis based on the binding strength of RAD21 revealed that specific forms of non-B DNA-forming sequence motifs were enriched in high- and low-binding subsets of RAD21-alone sites (Fig S3J). Together, this suggests that the increased ChIP-Seq signal of MRE11 and STN1 at the RAD21-alone sites could be explained by their underlying DNA sequence features that are susceptible to replication stalling and the cohesin complex could be recruited there to aid in DNA repair. Whereas at CBSs, the replication stalling could be influenced by the co-binding of CTCF and cohesin, along with the underlying DNA sequence features.

### Enrichment of DNA double-strand breaks at CTCF/cohesin-binding sites in the replication phase

Persistent replication stress leads to replication fork collapse and DSBs[35,36]. To check whether the CBSs undergo DSBs, we measured the enrichment of γH2AX (phosphorylation of Histone variant H2AX)[37,38], a known marker associated with DSBs, through ChIP-seq in the Mid S phase HeLa cells. We also measured the levels of H2AX in the Mid S phase (through ChIP-seq) as a baseline control. An overall enrichment of γH2AX (normalised to H2AX signal) at positions around CBSs was observed as compared to control sites (CTCF unbound and random sites) (Figs 3A and S4A). A similar signal of γH2AX was observed at RAD21-alone sites (Fig 3C) suggesting DSBs at RAD21 binding sites. Further analysis demonstrated that the increased γH2AX enrichment observed at CBSs is mainly from the regions with high CTCF and RAD21 co-binding (Fig 3E).

**Fig 3.**
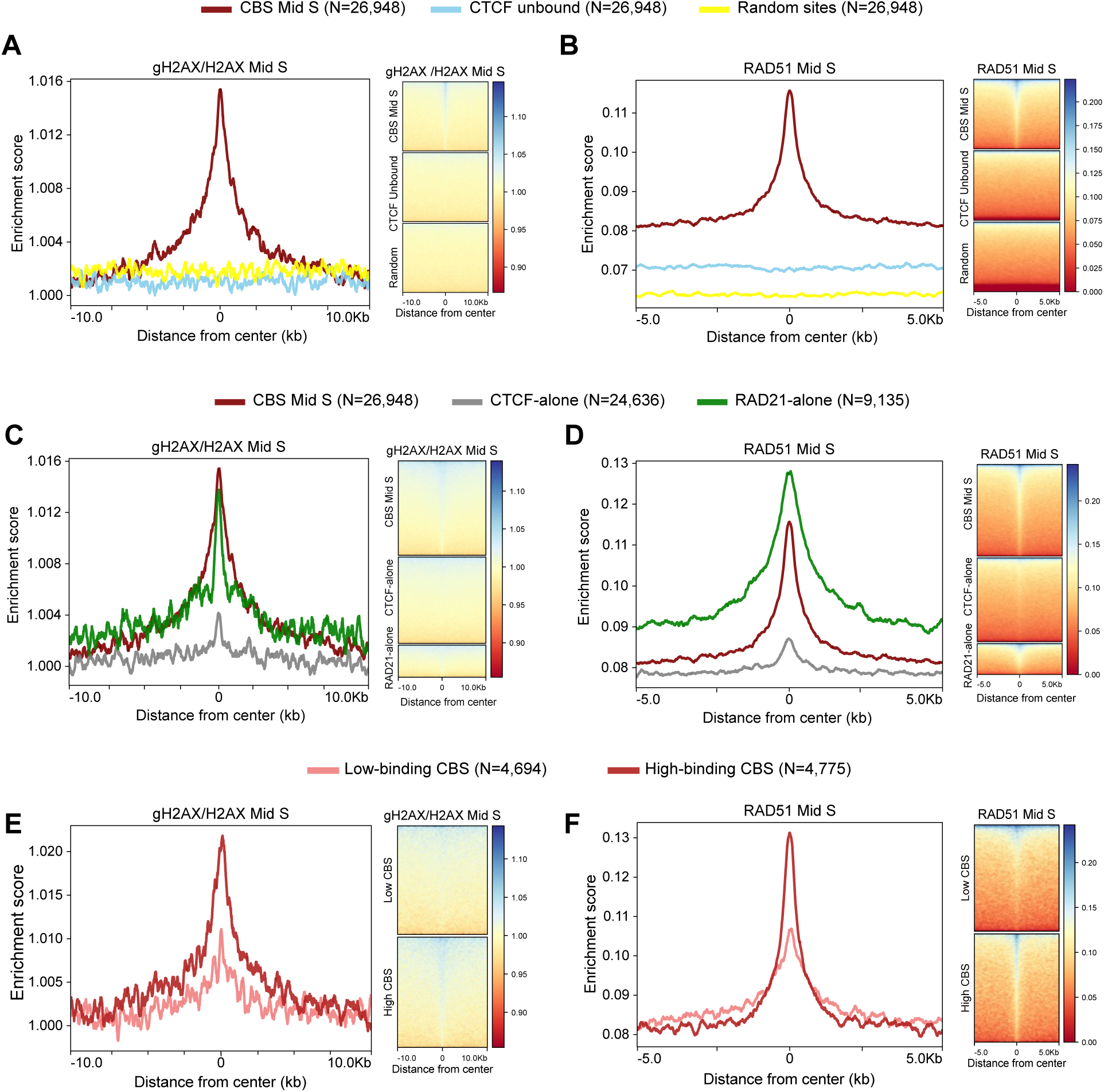
Enrichment of DNA damage response and repair proteins at CBSs. **A,B**: ChIP-seq signal profile and heatmap of **(A)** γH2AX (normalised with H2AX) and **(B)** RAD51 from Mid S HeLa cells plotted at CBSs (26,948 sites), CTCF unbound (26,948 sites) and random regions (26,948 sites). **C,D:** ChIP-seq signal of **(C)** γH2AX (normalised with H2AX) and **(D)** RAD51 plotted at CBS Mid S (26,948 sites), CTCF-alone (24,636 sites) and RAD21-alone (9,135 sites). **E,F:** ChIP-seq signal of **(E)** γH2AX (normalised with H2AX) and **(F)** RAD51 at low (4,694 sites) and high CBSs (4,775 sites). +/-10kb flanks were considered for the γH2AX signal, while RAD51 signal is plotted at +/-5kb regions.

Broken replication forks are repaired by Homologous recombination (HR) repair[39,40]. BRCA1 and RAD51 are the two main players in HR repair whose signals were found to be enriched at CBSs in asynchronous cells by re-analysing respective ChIP-seq datasets from ENCODE (Figs S4B-S4E). We asked if the HR repair proteins are recruited at CBSs in the S phase. For this, we performed a ChIP-seq of RAD51 in the Mid S phase HeLa cells. Strikingly, our ChIP-seq profile of RAD51 in the S phase displayed a higher signal at CBSs relative to its flanking regions as well as control sites (Figs 3B and S4F). In addition, the RAD21-alone sites also showed high enrichment of RAD51 (similar to MRE11 and γH2AX) (Fig 3D), suggesting a preferential repair at these sites. Further, the enrichment of RAD51 signals at CBSs positively correlates with CTCF/RAD21 binding strength at these sites (Fig 3F).

As DSB marker and repair proteins are highly enriched at CBSs, we tested the occupancy of the central DSB sensor kinase, ATM at CBSs in the Mid S phase of HeLa cells. The ChIP-Seq signal of the ATM also showed a higher enrichment at CBSs compared to the control sites (Fig S4G) and CTCF-alone sites (Fig S4H), and the enrichment at CBSs is associated with the binding strength of both CTCF and cohesin (Fig S4I). Moreover, the enrichment of ChIP-Seq signals of above proteins (γH2AX, RAD51 and ATM) at CBSs was relative higher than the DHS (open-chromatin regions), suggesting the enrichment is not biased by the chromatin accessibility (Figs S4J-S4L).

Together, our results support that the co-binding of CTCF/cohesin imposes replication stress, and CBSs are vulnerable to DNA damage. As a result, the DNA damage response/ repair (DDR) proteins are recruited to CBSs likely to prevent genomic instability.

### Exogenous replication stress increases DNA damage at CTCF/cohesin binding sites

Next, we asked whether the external replication stress could increase DNA damage or strand breaks at CTCF/cohesin sites during replication. For this, we performed ChIP-seq of DDR proteins in drug-induced replication stress conditions in HeLa cells through HU (which causes fork stalling) and Etoposide (ETO, which inhibits the re-ligation activity of topoisomerase II) treatments, separately. Overall, the enrichment of DDR proteins upon HU treatment was similar to the above results which were without exogenous stress in the Mid S phase (Figs S5). γH2AX signal at CBSs and other regions followed the same pattern after the induction of replication stress by 2 mM HU (Figs S5A and S5B). γH2AX enrichment was more prominent in prolonged HU treatment (16 hours) compared to that in the Mid S phase for 3 hours. MRE11, FANCD2 and ATM also displayed a higher signal at CBSs compared to control sites in HU-treated cells (Figs S5D, S5E, S5I and S5J). However, we observed a decrease in RAD51 binding at CBSs on HU treatment (Fig S5G) likely due to the persistent DNA damage and slower or impaired repair. Upon ETO treatment, Mid S HeLa cells also showed an enrichment of DNA breaks at CBSs (as seen from the γH2AX signal; Fig S5C) similar to previous findings in asynchronous cells[14,15,41]. However, MRE11 and RAD51 did not show any prominent localisation at CBSs (Figs S5F and S5H), possibly due to the perturbed DNA damage sensing and repair.

### Replication stress and DNA damage response at CTCF/cohesin-binding sites in normal cells

Since all the above experiments were performed in the cancerous cell line, we asked whether the replication stress-associated proteins are enriched at CBSs in normal conditions as well. For this, we first asked whether the CBS sites that are persistent (or common) between HeLa and normal cell lines show similar DNA damage response. To test this, we considered the following normal cell lines: RPE-1 (retinal pigment epithelial cells), IMR90 (normal lung fibroblast cell line) and H1 (human embryonic stem cells), for which the CTCF and cohesin (RAD21/SMC1) data was previously available to define CBSs (see Methods). Of the 26,948 CBSs in HeLa Mid S, 12,517 (46%) overlapped with CBSs of all three normal cell lines (referred as normal-shared CBSs). We plotted the DNA damage response signals from HeLa Mid S at these normal-shared CBSs as well as the CBSs specific to HeLa or normal cell lines. This showed that the normal-shared CBSs have high DNA damage response and DNA repair (MRE11, STN1, gH2AX, and RAD51) signal at the CBSs center as compared to immediate flanking region (Figs 4A-4D). However, this signal is relatively lower than HeLa-specific CBSs, but higher than the normal-specific CBSs, likely due to the cell-type specificity.

**Fig 4.**
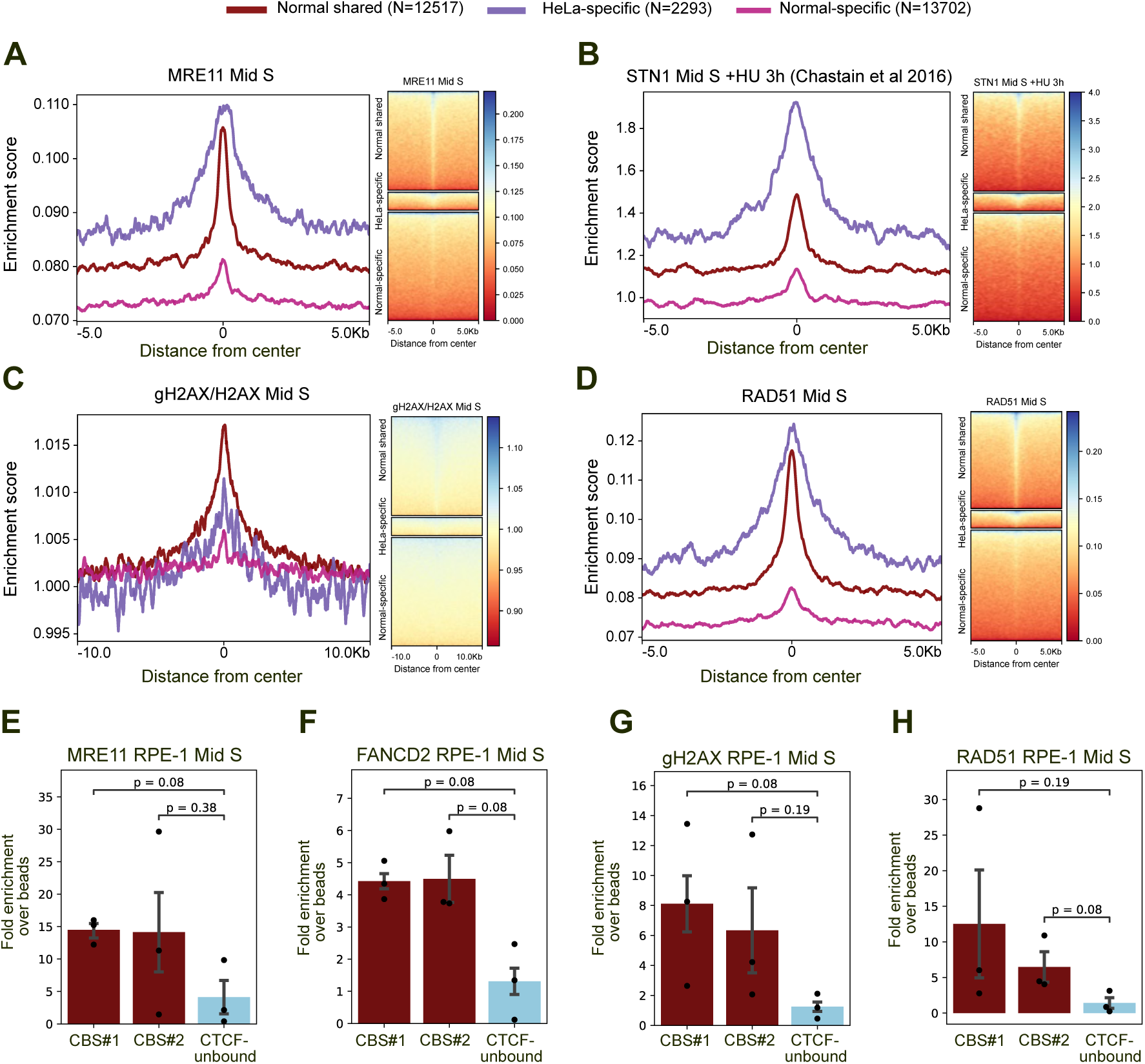
Replication stress and DNA damage response at CTCF/Cohesin-bindings sites in normal cells. **A-D**: ChIP-Seq profile of **(A)** MRE11, **(B)** STN1+HU 3h, **(C)** gH2AX(normalised with H2AX), **(D)** RAD51 in the Mid S phase of HeLa cells plotted at CBSs shared between HeLa and normal cells (H1, IMR90 and RPE1 cells) (12,517 sites) and HeLa-specific CBSs (2293 sites) and normal-specific sites (13,702). **E-H:** ChIP-qPCR plots of **(E)** MRE11 **(F)** FANCD2 **(G)** gH2AX and **(H)** RAD51 in Mid S synchronised hTERT RPE-1 cells at CBSs and CTCF-unbound sites. The y-axis (fold enrichment over beads) indicates the % input in immunoprecipitation (IP) divided by that of beads. Statistical significance is determined by a two-sided Mann-Whitney U test and error bars denote one standard error.

To further confirm this observation, we performed ChIP for MRE11, FANCD2, gH2AX and RAD51 in Mid S synchronised hTERT RPE-1 cells followed by qPCR at a few selected CBSs and CTCF-unbound sites (as a control) (Figs 4E-4H and Figs S6A-S6B, see Methods). Interestingly, we observed relatively higher enrichment of these proteins at CBSs compared to CTCF-unbound sites, suggesting that CBSs undergo replication stress/stalling even in normal cells. To check whether the removal of cohesin at these sites can reduce the replication stress or DNA damage, we performed RAD21 knockdown using siRNA(Fig S6C) followed by ChIP-qPCR for gH2AX and RAD51 proteins (Figs S6D and S6E). Although the DNA damage was lower at these CBSs sites, the signal was not stronger as compared to the control sites, and this is likely due to the global changes in the chromatin structure or additional DNA damages induced upon the RAD21 knockdown[42]. Further studies are required to explore this in detail. Taken together, this result suggests that many of the CBSs are persistent across cell-types, and thus the replication stress and DNA response observed at the CBSs are also seen persistent in cancer and normal cell lines.

### Increased somatic mutations at CTCF/cohesin-binding sites in tumours with loss of MRE11/STN1 genes

The association of DNA damage response proteins (MRE11 and STN1) with CTCF/ cohesin (Fig 2) in the S phase indicates that these proteins might be protecting CBSs from replication stalling and genomic instability. We wondered whether the rate of somatic mutations at CBSs would increase upon perturbations in MRE11 and STN1 (gene names MRE11A and OBFC1, respectively). Therefore, we explored somatic mutations at CBSs in patient tumours harbouring somatic copy number deletion in MRE11 and/or STN1 from the PCAWG[19] cohort. We computed the somatic substitutions at +/-1kb region centred around CBSs, CTCF unbound sites, CTCF-alone sites and RAD21-alone sites in tumour types with at least four samples in the following conditions: a) both MRE11 and STN1 wild-type (no alterations), b) either MRE11 or STN1 deletion, and c) both MRE11 and STN1 deletions. To account for the influence of DNA sequence composition on the mutation rate, we computed the expected mutation frequency based on the probabilities of each tri-nucleotide to be mutated (specific to each sample group) and calculated mutation fold change (fc: ratio of observed mutations by expected mutations at the centre +/-20bp) (see Methods). This showed that the enrichment of somatic mutation at CBSs in wild-type samples appeared to be high compared to flanking regions and control sites (Fig 5A), as shown previously[10–12]. Strikingly, we noticed that the tumours with deletions in MRE11-STN1 displayed a greater incidence of mutations at CBSs than wild-type samples in certain tumours (stomach, esophageal, liver, and breast cancers; Figs 5B and 5C).

**Fig 5:**
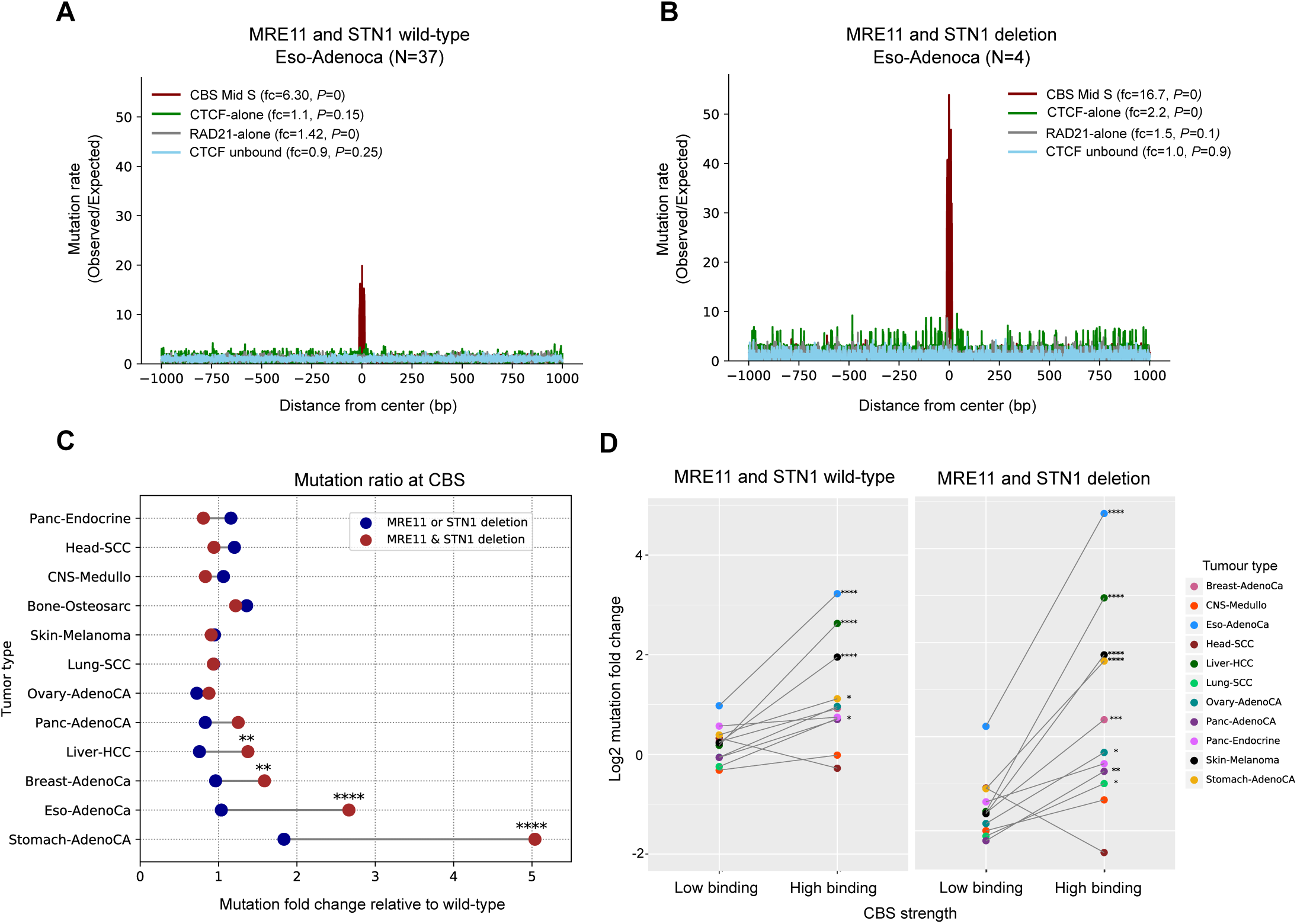
Enrichment of somatic mutations at CBSs in STN1/MRE11 deleted tumours. **A, B**: Mutation rate plotted at +/-1kb region of CBSs Mid S and control sites (CTCF unbound, CTCF-alone and RAD21-alone sites) in esophageal adenocarcinoma samples with **(A)** both MRE11 and STN1 wild-type (No. of samples, N=37) and **(B)** both MRE11 and STN1 deletion (No. of samples, N=4). P-values are calculated by using G-test. **(C)** Mutation fold change at CBSs (+/-20bp) core in tumour samples with either MRE11 or STN1 deletion (or both deletion) relative to mutation fold change in MRE11 and STN1 wild-type samples across different tumour types. **(D)** Mutation fold change at CBSs (+/-20bp) core stratified based on the CTCF and RAD21 binding strength. On the x-axis, the high binding represents CBSs with both CTCF and RAD21 strong binding and low binding represents CBSs with both CTCF and RAD21 weak binding. The statistical difference was calculated by using Fisher’s exact test. p-value annotation legend *: 0.01 < P <= 0.05, **: 0.001 < P <= 0.01, ***: P< 0.001, ****: P< 0.0001.

For instance, in esophageal adenocarcinoma, samples with deletions in both MRE11 and STN1 genes exhibited a dramatic increase (fc=16.7, P=0) in somatic substitution rate compared to MRE11-STN1 wild-type samples (fc=6.3, P=0) (Figs 5A and 5B). Interestingly, neither the MRE11-STN1 wild-type or deletion samples showed any increase in mutations at CTCF-unbound sites (both wild-type: fc=0.93, P=0.25; both deletion: fc=1.01, P=0.895) or RAD21-alone sites (both wild-type: fc=1.42, P=0; both deletion: fc=1.52, P=0.068). CTCF-alone sites and other transcription factor binding sites (TFBS) displayed a slight increase in mutation rate in MRE11 and STN1 deletion tumours (Fig 5B, Figs S7A and S7B), suggesting that the mutations are accumulated specifically at CBSs upon MRE11 and STN1 gene deletion. Moreover, the increase in mutation rate at CBSs was relatively higher in samples with both MRE11 and STN1 deletion as compared to samples with either MRE11 or STN1 deletion and sample with both wild-type (Fig 5C), especially in liver, breast, esophageal and stomach cancers (Fisher exact test, P <= 0.01). For example, stomach adenocarcinoma samples with either MRE11 or STN1 deletion (fc=3.41, P=0) exhibited a 1.8-times increase in mutations, while samples having both MRE11 and STN1 loss (fc=9.37, P=0) showed a 5-times increase in mutations at CBSs compared to MRE11 and STN1 wild-type samples (fc=1.86, P=0) of the same tumour type (Fig 5C). In contrast, mutations at control sites in MRE11-STN1 deletions were similar to that of wild-type samples in most tumours (Fig S7C).

As enrichment of DDR proteins at CBSs was associated with CTCF/RAD21 binding strength (Figs 2 and 3), we wondered whether the mutation rate at CBSs also follows the same trend. As expected, the high CBSs displayed an increased mutation rate compared to low CBSs across tumours (Fig S7D). Analysing mutations at low and high CBSs in MRE11-STN1 wild-type and deletion samples revealed a drastic increase in mutation rate at high CBSs in MRE11-STN1 deletion samples in many tumours suggesting the functional significance of DNA damage response proteins enrichment at CBSs (Fig 5D). Together, this suggests that the sequence compositions or binding of CTCF or RAD21-alone do not determine the elevated mutation rate at CBSs. Instead, the co-binding of CTCF and RAD21 contributes to a high mutation rate at CBSs and the mutation rate is strongly associated with their binding strength.

Mutation in the TP53 gene was previously associated with an increased somatic mutation rate at CBSs in colorectal cancers[12]. We wondered whether the increased mutation rate in esophageal cancer is influenced by TP53 mutation status. Towards this, we checked the mutations in samples having TP53 deletion ensuring that those samples do not have any alterations in MRE11 or STN1 genes (i.e. MRE11 and STN1 wild-type). Interestingly, the eso-adenocarcinoma samples with TP53 deletion did not show any increase in mutations at CBSs, suggesting that irrespective of the TP53 gene status, MRE11 and STN1 deletion leads to accumulation of mutations at CBSs in esophageal cancer (Figs S7E and S7F). However, we noticed biliary adenocarcinoma and stomach adenocarcinoma displaying high mutations at CBSs in samples with TP53 deletion (Fig S7G). We further explored these samples to see whether they harbour heterozygous (shallow deletion) or homozygous (deep deletion) in TP53 deletion. Interestingly, samples with homozygous TP53 deletion exhibited a higher mutation rate at CBSs compared to heterozygous condition (Figs S7H-S7J). Together, this suggests that the perturbation in DNA damage sensing/response genes could lead to increased somatic mutation rate at CBSs, because these sites experience high levels of DNA damage/DSBs.

## Discussion

CTCF and cohesin are involved in diverse functions like genome folding, transcriptional regulation[7],[43], DNA replication[44,45], DNA damage response[46] and repair[47]-[48]. Despite these roles in the maintenance of genomic integrity, the local DNA regions bound by both CTCF and cohesin (CBS) are found to be a hotspot for genomic instability[12,14,15],[41] and somatic mutations in multiple cancers[10–12],[49]. In particular, the CBSs at the TAD boundaries and late replicating regions showed high somatic mutations[13]. In some cases, mutations at TAD boundaries have been shown to disrupt TAD insulation, leading to the activation of proto-oncogenes[50]. However, the molecular mechanism underlying high somatic mutation rates at the CBSs remains unclear. Here, we show that the CBSs experience a higher amount of replication stress compared to the immediate neighbouring sites. This is associated with increased DNA double-strand breaks and enrichment of somatic mutations in cancer genomes.

Though the binding of CTCF and cohesin to chromatin are extensively studied in the G1 phase and mitosis[5,25,51–55], the presence of these proteins during the S phase and the hindrance posed by them to DNA replication are not yet fully understood. We find that the CTCF and cohesin proteins are present in chromatin-bound nuclear fraction throughout the cell cycle (Fig 1B). FRAP experiments revealed that the dynamics of CTCF present in the nucleus during S phase are similar to that of asynchronous cells (Figs 1C and 1D). This result is in line with the CTCF-FRAP experiment by Hansen et al[56] in mESCs. However, the cell cycle synchronisation method in our study ensures the selection of cells undergoing DNA replication, as compared to the aforementioned study where the S and G2 cells cannot be distinguished (due to the FUCCI system). Further, to map the genome-wide binding of CTCF and RAD21 in the S phase, we generated ChIP-seq data in synchronised HeLa cells. This revealed that most of the CTCF and RAD21 binding sites in G1, including the persistent or constitutively bound CTCF/cohesin sites in all cell types[57],[58], are occupied in the S phase too, providing independent validation of CTCF/cohesin binding in the replication phase. This is consistent with the previous observations based on HiC analysis that the global chromatin architecture is largely unchanged across interphase[59].

Next, we tracked the genome-wide binding of replication stress and DNA repair-associated proteins in the S phase to monitor endogenous stress at CBSs. Previous studies that performed ChIP-seq of DDR proteins were mainly in the exogenous stress conditions or enzyme-induced break scenarios. For instance, Chastain et al[32] uncovered that the STN1 is recruited to GC-rich fragile genomic regions upon replication stress (induced by exogenous HU treatment) to promote fork progression. Given that CTCF binding sites have a high GC content, we utilised their STN1 ChIP-seq dataset and observed that not only STN1 signal is high at CBSs but also correlated with the binding strength of CTCF/cohesin, suggesting that the CTCF/cohesin binding cause replication fork stalling. Complementing this, our MRE11 and FANCD2 ChIP-seq data also showed a high enrichment at CBSs in the replication phase with and without any exogenous damage, confirming replication stress/fork stalling at these sites (Fig 2). This high enrichment of MRE11 at CBSs could be partly explained by the physical interaction between CTCF and MRE11 at DNA damage sites as shown previously by Hwang et al[47]. Further, our γH2AX ChIP-seq data revealed persistent endogenous DSBs during replication at CTCF/cohesin binding sites. Although γH2AX signals enrich differently at various genomic sites depending on the kind of exogenous replication stress[60], we find a consistent occupancy of γH2AX at CBSs upon external replication stress induced by HU and ETO. In addition, we observed that the enrichment of DNA damage response/ repair protein is higher in RAD21-alone sites, followed by CBSs (both CTCF and cohesin binding) and CTCF-alone sites, suggesting that the binding of RAD21 is associated with replication fork stalling and DNA repair. This complements the previous finding that cohesin (RAD21) binding is associated with TOP2B-mediated DNA double-strand breaks at chromatin-loop anchors in asynchronous cells[14,15,61]. Also, the recent study by Peripolli et al[17] showed that increase in cohesin-loading, through oncogenic MYC induction, cause DNA replication stress in a CTCF-dependent manner in both normal and cancer cell lines at the global scale. Our study further compliments these findings by probing replication stress and DNA damage response specifically at the CTCF/cohesin binding sites as these sites exhibit unique features, both structurally and architecturally.

Finally, we analysed the essentiality of replication stress-associated proteins at CBSs by assessing the somatic mutation rates at CBSs in patient tumours harbouring MRE11 and STN1 gene deletion or not (wild-type) from the PCAWG[19] cohort. We found that MRE11 and STN1 deletion is associated with increased mutation rates at CBSs in certain tumours (stomach, esophageal, liver and breast cancers), suggesting that this mechanism is specific to those cell types. Loss of DNA damage sensors like MRE11 and STN1 leads to inefficient replication stress response and repair that increases the likelihood of mutation rate across the genome. However, the control sites used in this study (CTCF unbound, CTCF-alone and RAD21-alone sites) did not show any drastic increase in mutations relative to CBSs in MRE11-STN1 deletion samples even after correcting for background mutations, highlighting the requirement of CTCF and cohesin together to cause genomic instability. This further supports the previous findings which showed that the enrichment of somatic mutations at CBSs is relatively higher as compared to CTCF- or RAD21-alone sites[12] and the recurrent mutation at CBSs is associated with chromosomal instability in the nearby regions[49].

During replication, DNA-binding proteins ahead of the fork are temporarily removed from the DNA to ensure the proper propagation of the replication fork[3,62–65]. Recent studies [66,67] have shown that the CTCF can be bound back to its motif sites post-replication to facilitate the nucleosomal position in the neighbouring regions. On the other hand, the binding of transcription factors to DNA damage has been shown to impair DNA repair activity[10,68–70]. Based on these, we speculate that at CBSs (i.e., CTCFcohesin co-bound sites), the rapid re-binding of CTCF could impair the repair of errors (DNA mismatches) introduced during DNA replication, resulting in the accumulation of mutations at these sites, particularly in MRE11/STN1 deleted conditions (Fig 6). At CTCF-alone sites, this was not the case due to less replication stress and DSB observed at these sites and this can reduce the chance of DNA mismatches at these sites. Though the RAD1-alone sites showed high replication stress, the higher repair activity observed at these sites (likely due to the involvement of cohesin in DNA repair activity as well) could probably reduced the source of DNA mismatches or the nature of cohesin binding (non-sequence-specific) not affected the DNA repair activity at these sites.

**Fig 6:**
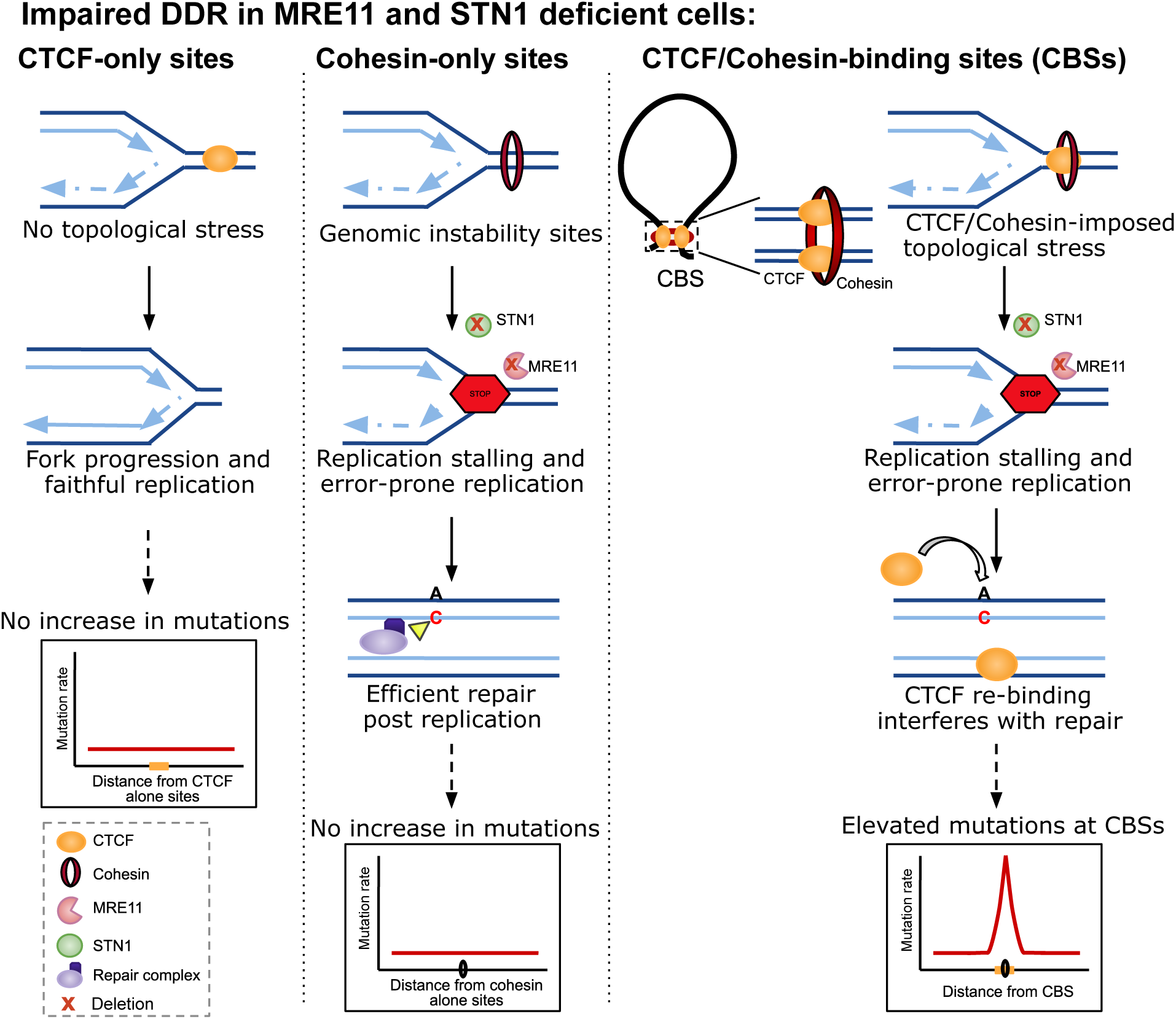
Schematic model for the possible mechanism underlying elevated somatic mutations at CBSs in cancers upon MRE11 and STN1 deletion. At CTCF-only bound sites, the replication fork progresses without any constraints and undergoes faithful replication, thus preventing the enrichment of somatic mutations at these sites. Whereas cohesin-only bound sites are enriched with DNA sequence susceptible to genomic instability and replication stalling. This can lead to error-prone replication; however, faithful repair post-replication prevents the enrichment of somatic mutations at these sites. However, at CTCF/cohesin-binding sites (CBSs), CTCF and cohesin co-binding induced stress can lead to replication stalling and error-prone replication. Moreover, the immediate re-occupancy of CTCF post-replication at these sites could interfere with mismatch recognition and repair, leading to an elevated mutation rate at CBSs.

However, the limitations of the study include: a) the DNA replication fork stalling and DNA strand breaks were measured indirectly through the ChIP-seq of DNA damage sensing/repair proteins, but this can be further improved with proximity ligation assay involving replication complex protein and cohesin, or with replication dynamics assays specifically focused on CBSs; b) the overall ChIP-seq signal of above DNA damage sensing/repair proteins was weak, as compared to CTCF or RAD21. This is likely due to the non sequence-specificity and transient binding of the DNA repair proteins. However, despite this weak signal, we observed a strong enrichment at CBSs, as compared to control regions; c) given that the genome organisation and chromatin loops are correlated with the replicating timing[44], the DNA strand breaks (and enrichment of histone marks associated with that) around CBSs could be partly associated with the replication initiation; d) whether the activity of error-prone replication (e.g., break-induced replication) or error-prone repair (e.g., translesion synthesis) that underlies the source of DNA mismatch at CBSs remains to be characterised; e) the somatic mutations analysed here were predominantly from primary tumours, thus extending our analysis to normal tissues could help to understand the baseline rate of these events, and to metastatic tumours (which undergone topoisomerase inhibitor or other chemotherapy treatment) to study the impact of treatment induced stress at CBSs.

In summary, our study demonstrates that binding of the CTCF/cohesin on DNA imposes constraints on the DNA that affect the progression of replication fork, leading to fork stalling and DSBs. This could be a potential source of mutations at CBSs, and the likelihood of this is higher in cells lacking DNA damage sensing/repair proteins (such as MRE11 and STN1). This finding has important implications in understanding how chromatin architecture influences the somatic mutational processes across the genome and also for the development of better background mutation models to predict non-coding cancer drivers.

## Materials and Methods

### Cell lines and culture conditions

HeLa cells (obtained from ATCC) were grown in High Glucose Dulbecco’s modified Eagle’s medium (DMEM) (Gibco, ThermoFisher, #10569044), supplemented with 10% Fetal Bovine Serum (FBS, Gibco, Invitrogen #16000044) and 1% penicillin-streptomycin (Gibco, Invitrogen #15140163). hTERT RPE-1 cells were cultured in DMEM/F12 (Gibco, Thermofisher #10565042) with 10% FBS and 1% penicillin-streptomycin. Cells were cultured at 37°C in a humidified 5% CO_2_ incubator and passaged every 3-4 days before reaching confluency. Cells were routinely tested for free of mycoplasma by using LookOut Mycoplasma PCR detection kit (Sigma, #MP0035).

### Cell cycle synchronisation

Cell cycle synchronisation was achieved following widely used thymidine arrest/release or nocodazole block method[71]. Briefly, HeLa cells were cultured at 60-70% confluency and treated with 2 mM Thymidine (Sigma #T1895) for 16 hours. Cells were released from the Thymidine block by washing twice with PBS and incubating in fresh DMEM (as mentioned above ) for either 3 (Early S), 4 (Mid S), or 5 (Late S) hours, before harvesting. G1/S cells were obtained by collecting cells after the Thymidine block without release. To obtain mitotic cells, 200µg/ml of Nocodazole (Sigma, #M1404) was added to the culturing media and incubated for 24 hours. Cell cycle synchronisation was confirmed by the flow cytometry profile. Replication stress was induced by treating Mid S synchronised cells with 2 mM HU (Sigma, #H8627) or 25µM etoposide (Sigma, #E1383) for 3 hours. For prolonged HU treatment, cells were incubated with 2 mM HU overnight. The same protocol was followed for synchronising hTERT RPE-1 cells.

### Flow cytometry

Synchronised HeLa cells were trypsinized and collected in DMEM. The cell pellet was washed twice with ice-cold PBS, resuspended in DMEM and fixed with 80% ethanol with constant vortexing, followed by incubation on ice for 10-15 minutes. Fixed cells were washed with PBS, stained with 50µg/ml Propidium Iodide (Invitrogen, #P3566) containing RNase A (Qiagen, #19101), incubated at RT for 30 minutes, and measured the DNA content using a flow cytometer (BD FACSVerse) at CIFF Facility, NCBS. The FCS files obtained for DNA content profiles were analysed for the proportion of cells in each cell cycle stage using the Watson Pragmatic model[72] in FlowJo^TM^ Software (BD Life Sciences). The same protocol was followed for hTERT RPE-1 cells.

### Immunostaining

HeLa cells were seeded at a density of 0.3x10^5^ cells on coverslips and synchronised to the Mid S phase as described above. Cells were then fixed using pre-chilled methanol at room temperature (RT) for 7 minutes, followed by three PBS washes. Cells were then incubated with a permeabilization buffer: 0.5% Triton X-100 (Sigma, #T8787) in 1% BSA (Invitrogen, #AM2616) at RT for 15 minutes. Next, primary antibodies (CTCF: D31H2 CST 1:600, MRE11: ab214 Abcam 1:600, PCNA: ab29 Abcam 1:500, RAD21: ab992 Abcam 1:500, STN1: sc-376450 1:200) were added in a 1% BSA in PBS solution overnight at 4°C or RT for 3 hours. Following primary antibody incubation, cells were washed with PBS-T (PBS + 0.1% Tween 20) three times, each of 5 minutes. Next, cells were incubated with a secondary antibody (Anti-Rabbit Alexa Fluor-568: Invitrogen A10042; Anti-Mouse Alexa Fluor-488: Invitrogen #A11001) at 1:500 dilution in PBS-T for 1 hour at RT and then washed three times in PBS-T. Nuclei were stained with 0.5µg/ml DAPI (Sigma, #10236276001) for 5 minutes and coverslips were then mounted onto glass slides using 90% glycerol. Images were acquired using ZEISS LSM 980 confocal microscope at the CIFF Facility, NCBS. Images were acquired in 0.15µm intervals with a minimum of 5 stacks with Airyscan mode to obtain super-resolution images. Image analysis was performed using Fiji[73].

### Colocalization analysis

The region of interest (ROI) was generated around the nuclear boundary on the DAPI channel to get the nuclear signal from every single cell. Colocalization was quantified using the JACoP (Just another Colocalization Plugin)[74] plugin on Fiji[73]. Colocalization was quantified with two different approaches: Pearson’s correlation coefficient (PCC)[29] and Mander colocalization coefficient[30]. PCC measures the correlation between intensity variation in channel 1 compared to channel 2. Mander’s coefficients (M1 and M2; where M1 is the fraction of channel 1 overlapping with channel 2 and M2 is the fraction of Channel 2 overlapping with channel 1) quantify the co-occurrence of signals[75]. Threshold values for Mander’s coefficients (tM1 and tM2) Costes automatic threshold methods[76] and visually checked for appropriate adjustments.

A minimum of 30 cells (n) were analysed for colocalization per biological replicate (N). The results were verified by performing each experiment independently at least three times. A two-sided non-parametric Mann-Whitney U test was performed to check whether there is a difference in colocalization in asynchronous and Mid S phase cells. The plots were generated using the Seaborn package in Python.

### Fluorescence Recovery after photobleaching (FRAP)

HeLa cells cultured on coverglass dishes were transfected with pKS070-pCAGGS-3XFLAG-(human)CTCF eGFP, a gift from Elphege Nora (Addgene plasmid #1564488)[6], using Lipofectamine 2000 (Invitrogen, 11668030) and OptiMEM (Gibco, #11058021). Cells were synchronised to the S phase 24 hours post-transfection by thymidine block and released into phenol red-free media for 4 hours (live imaging starts from 3.30 hours of release and ends at 4.30 hours). FRAP experiments were performed on Olympus FV3000 6 laser microscope at the CIFF facility, NCBS. A rectangular region of interest (ROI) of ∼1µm area was bleached using a 488 nm laser with 100% power. Throughout the imaging, cells were maintained at 37°C and 5% CO2. The images were captured before bleaching (5 frames), during bleaching (5th frame) and post-bleaching (rest of the frames) every 3 seconds for 3 minutes. Fluorescence recovery in the ROI across the frames was calculated by compensating background intensity as well as the variations in intensities in the same nuclei using the FRAP Norm plugin in Fiji. Data were derived from three biological replicates. For plotting the curve, the intensity obtained from above was normalised with the whole-cell intensity. As a control, the intensity at an unbleached ROI of the same area was also calculated and plotted similarly. The half-time of recovery (t_1/2_) was calculated by fitting an exponential curve to the recovery data.

### Fractionation

Synchronised HeLa cells were washed twice with PBS, and spun down at 2900 rpm at 4°C for 5 min. To the pellet, Buffer A (100 mM NaCl, 300 mM Sucrose, 3 mM MgCl2, 10 mM PIPES pH 6.8, 1 mM EGTA and 0.2% Triton X-100) and 1x Protease Inhibitor Cocktail (PIC) were added and incubated on ice for 30 min with a gentle tap on the tube every 5 minutes. This was followed by centrifugation at 2900 rpm for 5 minutes at 4°C to get the cytoplasmic protein fraction in the supernatant. To the supernatant, a 6x Laemlli sample buffer (375 mM Tris-HCl pH 6.8, 9% SDS, 50% Glycerol, 0.03% Bromophenol blue and 9% beta-mercaptoethanol) was added and heated at 95°C for 15 minutes. The pellet was washed twice with Buffer A followed by incubation with Buffer B (1 M HEPES pH 7.6, 50% Glycerol, 3 M MgCl2, 0.5 M EDTA, 10% NP-40 and 2 M Potassium acetate) containing 1x PIC on ice for 30 minutes with a gentle tap on the tube every 5 minutes. The resulting solution was centrifuged at 12,000 rpm for 12 minutes and washed twice with Buffer B. To the pellet, which has insoluble chromatin-bound proteins, 2x Laemlli sample buffer was added and preheated at 95°C for 15 minutes.

### Western blotting

Cells were washed with PBS, scraped off the plate and then lysed and collected in a 2x Laemlli sample buffer. Cell lysates were pre-heated at 95°C for 5 minutes and spun down at 13,400 rpm for 1 minute at RT. Samples were separated on SDS-PAGE (stacking 4% and resolving (8% top and 15% bottom)) and the protein-resolved gel was transferred onto a PVDF membrane (Millipore, #IPVH00010) in a 1X transfer buffer (25 mM Tris, 192 mM glycine, pH 8.3 and 20% methanol) at 20 V for 1 hour. The membrane was blocked in 5% skimmed milk (Santa Cruz, sc-2324) prepared in 1x TBS-T at RT for 1 hour on the rocker. Blots were then incubated with primary antibody (Mouse anti-GAPDH Santa Cruz sc-47724; 1:6000, Rabbit anti-CTCF Cell Signalling Technology D31H2; 1:6000, Rabbit anti-RAD21 abcam ab992 1:2000, Rabbit anti-Histone H3 Sigma H0164; 1:6000, Mouse anti-Cyclin A Santa Cruz sc239- ; 1:2000, Mouse anti-Cyclin B1 Santa Cruz sc-245; 1:2000) overnight at 4°C on the rocker. After primary incubation, blots were washed thrice with 1x TBS-T for 10 minutes each and incubated with HRP conjugated secondary antibody (anti-Rabbit HRP Invitrogen, #32460; 1:10,000 or anti-Mouse HRP Invitrogen, #32430; 1:10,000) prepared in 1xTBS-T for 1 hour at RT on the rocker. Blots were further washed thrice with 1x TBS-T and developed using Clarity Western ECL (BioRad, #1705060). Images were captured using the ImageQuant LAS 4000 gel documentation system. Images were then processed using ImageJ software.

### Chromatin immunoprecipitation (ChIP)

Cells were cross-linked using 1% formaldehyde (Sigma, #F8775) for 10 minutes and then were quenched with 125 mM glycine (Sigma, #50046) for 5 minutes at 60 rpm at RT. Cells were washed thrice with ice-cold PBS, scraped and pelleted down at 2000 rpm at 4°C for 5 minutes. The cell pellet was either stored at -80°C or directly used in further steps. First, the cells were lysed with L1A buffer (10 mM HEPES/KOH pH 7.9, 95 mM KCl, 1 mM EDTA pH 8 and 1% NP-40) for 10 minutes on ice and centrifuge at 3500 rpm for 5 minutes. Nuclei were obtained by adding a nuclear lysis buffer (L2) (50 mM Tris-HCl pH 7.4, 1% SDS and 10 mM EDTA pH 8) supplemented with 1x PIC, to the pellet for 10 minutes on ice. The chromatin was fragmented to an average size of 400-500 bp by 22 cycles of 30 seconds ON and 30 seconds OFF pulses using a Diagenode Bioruptor sonicator. Fragment size was visualised by running the chromatin on 1% Agarose gel. The DNA concentration was checked in NanoDrop. Chromatin lysate was cleared by centrifugation at 12000 rpm at 4°C for 12 minutes. One hundred µg of chromatin lysate was diluted in the Dilution buffer (20 mM Tris-HCl pH 7.4, 100 mM NaCl, 2 mM EDTA pH 8 and 0.5% Triton X-100) in a 1:1.5 ratio and 10% of chromatin lysate was taken as input. The antibody was added and incubated overnight at 4°C with a constant rotation. Antibodies used are: CTCF (D31H2: Cell Signaling Technology, 1µg), RAD21 (ab992: Abcam, 1µg), MRE11 (ab208020: Abcam, 2µg), γH2AX (ab81299: Abcam, 2µg), H2AX (ab11175: Abcam, 2µg), ATM (ab201022: Abcam, 2µg), FANCD2 (NB100-180: Novus Biologicals, 2µg) and RAD51 (ab176458: Abcam, 2µg).

The next day, Protein A/G Dynabeads (Invitrogen, #10001D and #10003D) which were blocked in 1% BSA (Invitrogen, #AM2616) for 1 hour, were added (15µl per tube) for 4 hours at 4°C at 12 rpm. This was followed by washes with wash buffer I (20 mM Tris-HCl pH 7.4, 150 mM NaCl, 0.1% SDS, 2 mM EDTA pH 8 and 1% Triton X-100), wash buffer II (20 mM Tris-HCl pH 7.4, 500 mM NaCl, 2 mM EDTA pH 8 and 1% Triton X-100), wash buffer III (20 mM Tris-HCl pH 7.4, 250 mM LiCl, 1% NP40, 1 mM EDTA pH 8 and 10% Sodium deoxycholate) and 1xTE buffer (10 mM Tris pH 8, 1 mM EDTA pH 8) in sequence for 5 minutes at 4°C. Flow through was discarded after each wash step. Then the protein-DNA complex was eluted in 200µl of Elution Buffer (100 mM NaHCO3, 1% SDS) at 37°C. Decrosslinking was performed by adding 14 µl of 5M NaCl (Sigma, #746398) to the eluate at 65°C overnight. The DNA was purified using the Phenol:Chloroform:IAA (Invitrogen, #AM9732) method followed by ethanol precipitation. Finally, the pellet was dissolved in water and the extracted DNA was used in a qPCR reaction to verify the ChIP and proceeded for sequencing.

### siRNA mediated knockdown

S M A R T P o o l s i R N A s a g a i n s t R A D 2 1 (#L-006832-00-0005) and scramble (#D-001810-10-05), were purchased from GE Dharmacon. Transfections were performed using Lipofectamine 2000 (Invitrogen #11668027). To increase the knockdown efficiency, two rounds of transfection were performed.

### ChIP-sequencing and analysis

ChIP sequencing was carried out at the NGS Facility, NCBS. Library preparation was done using the NEBNext^Ⓡ^ Ultra^TM^ II DNA Library preparation kit (NEB, #E7103L). Libraries were sequenced on single-end 50 bp (1x50 bp) reads on the HiSeq 2500 Illumina platform. ChIP-seq data were analysed using the nf-core/ChIPseq[77] pipelines. The command used was ‘nextflow run nf-core/ChIPseq -r 1.2.1 -profile docker --input design.csv --genome GRCh37 --single_end --narrow_peak’ for narrow peaks. Briefly, the pipeline trims the adapters (Trim Galore!), aligns the reads to reference genome (BWA), marks duplicates (picard), filters (SAMtools and BEDTools), performs alignment-level QC and library complexity estimation (picard, Preseq), creates normalised bigWig files (BEDTools, bedGraphToBigWig) and calls peaks (MACS2). For CTCF and RAD21 ChIP, MACS2 peak calling was run by specifying ‘narrowpeak’ while all other ChIP peaks were called by ‘broadpeak’. Overlapping ChIP-seq peaks between the biological replicates were computed with BEDTools[78] intersect by keeping the fraction of overlap as 0.5.

### ChIP-qPCR

A qPCR reaction was prepared in triplicates using the SYBR™ Green PCR Master Mix (Applied Biosystems, #4309155). The ΔΔCT method was used to analyze the qPCR data and calculate the % input value. This was further normalized with beads to get the fold enrichment over the beads.

### Somatic mutation analysis

Somatic mutation and somatic copy number alteration (SCNA, gene level) data of 37 tumour types were obtained from PCAWG[19]. Samples with mutations in the exonuclease domain of POLE were excluded from the analysis. Single base substitution mutations were extracted and intersected with each of the regions (i.e., CBSs, CTCF unbound sites, CTCF-alone sites, RAD21-alone sites and TFBS) spanning 1000 bp upstream and downstream (+/-1kb) from the motif/peak centre, using BEDTools intersect to compute the rate of substitutions at respective sites. The samples were grouped into three categories according to the SCNA values of MRE11 (MRE11A) and STN1 (OBFC1) genes (samples with SCNA value of zero for both MRE11A and OBFC1 was considered as wild-type or no alterations; and samples with SCNA value less than 0 was considered as deletion). Samples with amplifications (SCNA >0) in MRE11 or STN1 were not considered for the analysis. Also, tumour types with less than four samples in any of the groups were excluded. Similarly, samples with TP53 gene deletion were classified as above. Further, the samples with SCNA values -1 and -2 were categorised as likely heterozygous and homozygous deletion, respectively.

To assess whether the mutations observed at CBSs (or other TFBS) are due to the local DNA sequence contexts, we calculated the background mutations based on the probability of each trinucleotide to be mutated (as described in Frigola et al[79]). Briefly, for each sample group (wild-type or STN1-MRE11 deletion samples), we computed the mutation probability by using the frequency of mutations observed in the specific tri-nucleotide context normalised by the abundance of that tri-nucleotide context in the genome. Fold change (fc) or the enrichment of mutations over background was computed as the ratio of observed mutations over expected mutations at +/-20bp from midpoint (core region). The significance value was calculated by using G-test goodness-of-fit (frequency of observed mutations in core and flanking regions were compared to the frequency of expected mutations in those regions).

### Predicting CBS in the S phase

To identify the CTCF/cohesin binding sites in the S phase (CBSs Mid S), we first performed motif analysis in 87,734 CTCF peaks (from HeLa Mid S phase) to check the presence of CTCF motif using MOODS (Motif Occurrence Detection Suite)[80] in Python with a p-value cut-off of 0.0001. The position frequency matrix (PFM) file of CTCF motif in humans (Matrix ID: MA0139.1) was obtained from JASPAR[81,82]. In case of a peak region with multiple CTCF motif matches, the one with maximum motif match score was retained. Further, we defined CTCF/cohesin-binding sites (CBSs, n=26,948) as those CTCF peaks with CTCF motif match and have a RAD21 peak within 100 bp from the CTCF motif centre (calculated using BEDTools closest function). We separated the CBS Mid S into 3 quantiles; low, medium and high based on CTCF and RAD21 ChIP-seq signals. Those CBSs with both CTCF and RAD21 low binding strength were defined as Low CBS and both CTCF and RAD21 high binding strength were considered as High CBS. CBSs from other cell lines were also obtained by following the same method as above on the ChIP-seq data obtained from ENCODE (see “External datasets” section below).

Control sites were defined as follows: CTCF unbound sites were obtained by taking the CTCF prediction sites from JASPAR[81] and removing all overlapping CTCF ChIP-seq sites in the S phase. The origin of replication sites (from ORC1 ChIP-seq in HeLa[83]) and DHS were also removed from these regions and randomly sampled 26,948 sites to match the number of CBSs in the S phase. Random sites used in this study were the randomly sampled genomic regions of 26,948 sites with no CTCF/RAD21 ChIP enrichment or replication origin sites or DHS.

### Overlap with Non B forms of DNA and CFS

Overlap of various genomic sites (eg., CBSs) with Non-B forms and CFS were determined by intersecting their genomic coordinates using BEDTools[78] intersect and the fraction of genomic regions that overlapped was calculated as the sum of overlap regions divided by the total size of feature of interest (e.g., CBSs).

### ChIP enrichment analysis

We used deepTools computeMatrix to check the occupancy of proteins centred at specific genomic regions. ChIP-seq bigwig files were provided as the score files (-S) and genomic sites of interest in bed format as region files (-R). The enrichment was plotted by plotHeatmap and plotProfiles functions.

### External datasets

To define CBSs in other cell lines, CTCF and RAD21 ChIP-seq peaks from various human cell lines were downloaded from ENCODE[26] from the GEO: GSE51334, which includes HeLa (cervical cancer cell line), A549 (lung cancer cell line), H1 (human embryonic stem cells), HCT116 (colorectal cancer cell line), HepG2 (liver cancer cell line), IMR 90 (normal lung fibroblast cell line), MCF7 (breast adenocarcinoma cell line), RPE-1 (human retinal pigment cells) and SK-N-SH (human neuroblastoma cell line). SMC1 ChIP-Seq data was downloaded from Peripolli et al[17]. CBSs in LoVo (colorectal cancer cell line) were from Katainen et al 2015[12]. STN1 ChIP-seq data in HeLa cells treated with HU[32] were obtained from the GEO: GSE82123. The origin of replication sites in HeLa was taken from the ORC1 ChIP-seq data[83] from the GEO: GSM922790. Genomic annotation of Non-B forms of DNA was obtained from Cer et al[84]. Human chromosome fragile sites data was obtained from Kumar et al[85]. DHS sites were downloaded from ENCODE[26] and removed CTCF/RAD21 and origin overlapping sites. For analysis, 26,948 DHS sites were randomly sampled to match the number of CBSs.

### Statistics

Half-time recovery (t1/2) of CTCF FRAP in asynchronous cells was compared to Mid S cells using a two-sided Mann-Whitney U test. The p-values for protein colocalization were determined using a two-sided Mann-Whitney U test. The statistics for ChIP-qPCR data was determined by a two-sided Mann-Whitney U test. For mutation analysis centered at CBSs and control region, the p-value was determined by a G-test, where the likelihood ratio of observed versus expected mutations in the core 20 bp region is compared to the observed versus expected ratio in the 20bp flanking region. The mutation fold change at CBSs in MRE11 and/or STN1 deletion samples relative to MRE11 and STN1 wild-type samples was compared using Fisher’s exact test. The same test was also used to assess the significance of mutation fold change at control sites as well as to compare mutations at low versus high binding CBSs.

## Acknowledgements

We thank Aswin Sai Narain Seshasayee, Madhusudhan Srinivasan, and the members of RS and DN labs for feedback and suggestions on the manuscript. We acknowledge the Next Generation Genomics Facility (NGGF) and Central Imaging & Flow Cytometry Facility (CIFF) at NCBS for their support and help. The results shown here are in part based upon data generated by the TCGA Research Network: https://www.cancer.gov/tcga.

## Funding

This work was supported by the DBT/Wellcome Trust India Alliance Fellowship [grant number IA/I/20/1/504928] awarded to RS. We also acknowledge support of the Department of Atomic Energy, Government of India, under Project Identification No. RTI 4006 and intramural funds from NCBS-TIFR.

## Author contributions

EEF and RS conceived the study. EEF, DN and RS designed the experiments. EEF performed all the experiments and data analysis, interpreted the results, and prepared figures. DN and RS jointly supervised the study and interpreted the results. FEE and RS wrote the manuscript with inputs from DN. All authors read and approved the final manuscript.

## Data availability

The ChIP-seq data (fastq and processed files) generated from this study is available in the NCBI GEO (Accession ID: GSE292153).

## Supplementary Figures

**Fig S1.**
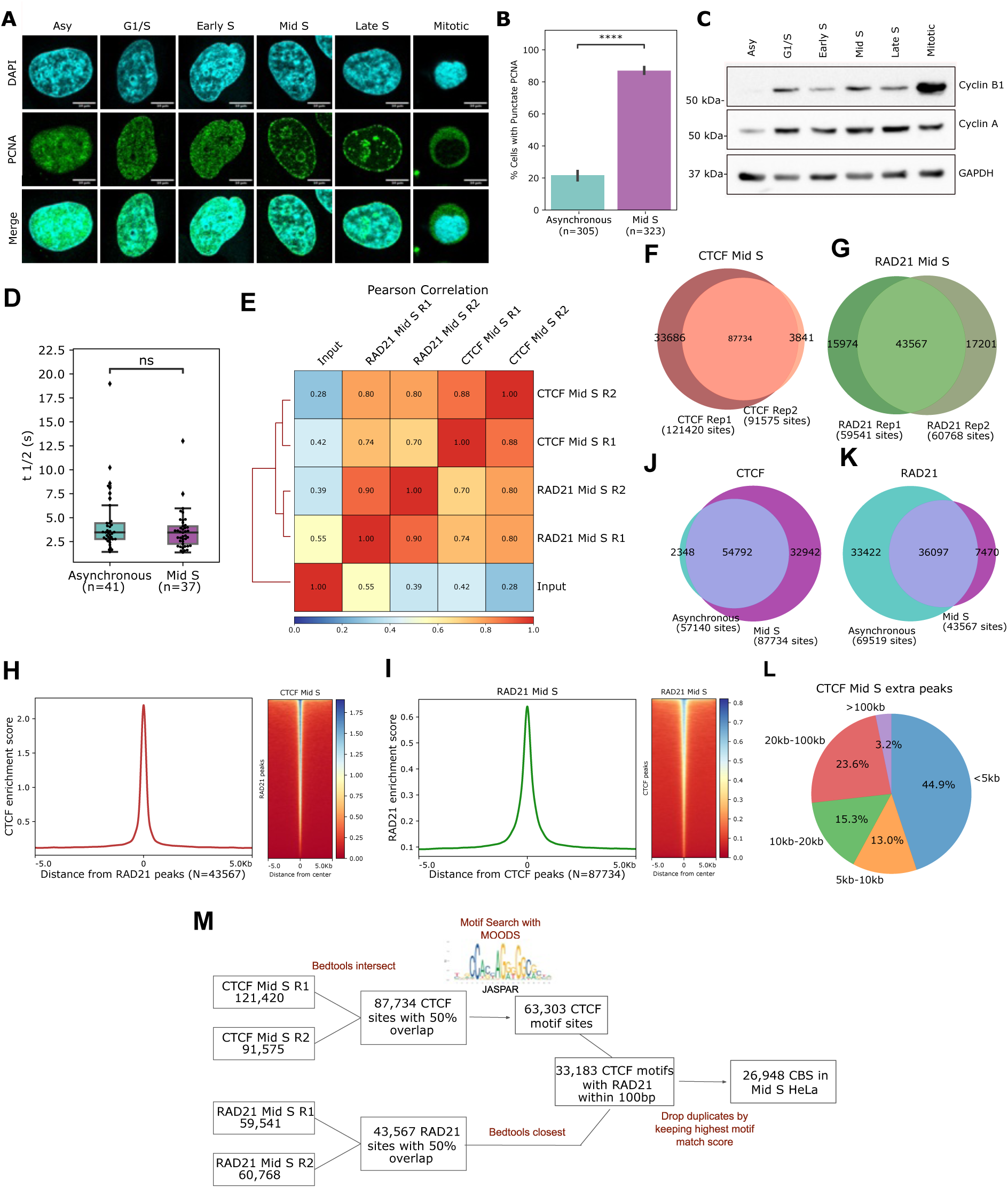
Profiling of CTCF/cohesin binding in replication phase. **(A)** Representative image of PCNA (in green) immunostained cells in various cell cycle phases. The nucleus is stained with DAPI (cyan). Scale bar 10µm. **(B)** Quantification of punctate PCNA in asynchronous and Mid S synchronised HeLa cells from five biologically independent experiments. The bar represents mean value and the error bar represents SD. The p-value was calculated using unpaired t-test. ****: p<00001. **(C)** Western blot showing expression of various Cyclin B and Cyclin A in the synchronized population of cells at different cell cycle phases. GAPDH was used as a loading control. **(D)** Half-time recovery (t1/2) of CTCF FRAP in asynchronous and Mid S HeLa cells. Centerline: median; Box limits: 1st and 3rd quartile; whiskers: maximum and minimum values: 1.5xIQR. The p-value was calculated using two-sided Mann-Whitney U test: ns: p>0.05. **(E)** Hierarchically clustered correlation heatmap of CTCF and RAD21 ChIP-seq replicates in Mid S HeLa cells. DeepTools’ multiBamSummary followed by plotCorrelation functions were used to find the Pearson correlation between the samples. **F, G:** Venn diagram showing the number of shared (with 50% overlap between the peaks) and unique peaks of **(F)** CTCF and **(G)** RAD21 biological replicates in the Mid S HeLa cells. **H, I:** ChIP-Seq coverage signals of **(H)** CTCF in Mid S synchronized HeLa cells plotted at ChIP-Seq peaks of RAD21 and **(I)** vice versa. **(J)** Intersection of CTCF ChIP-seq peaks in asynchronous and Mid S HeLa cells (see Methods). **(K)** Intersection of RAD21 ChIP-seq peak in asynchronous (from ENCODE) and Mid S HeLa cells. **(L)** Pie chart depicting the distance between additional CTCF ChIP-seq peaks observed in Mid S with the common peaks from Mid S and asynchronous cells. **(M)** Workflow for defining CTCF/cohesin binding sites (CBSs; i.e., regions bound by both CTCF and RAD21) in Mid S HeLa cells.

**Fig S2.**
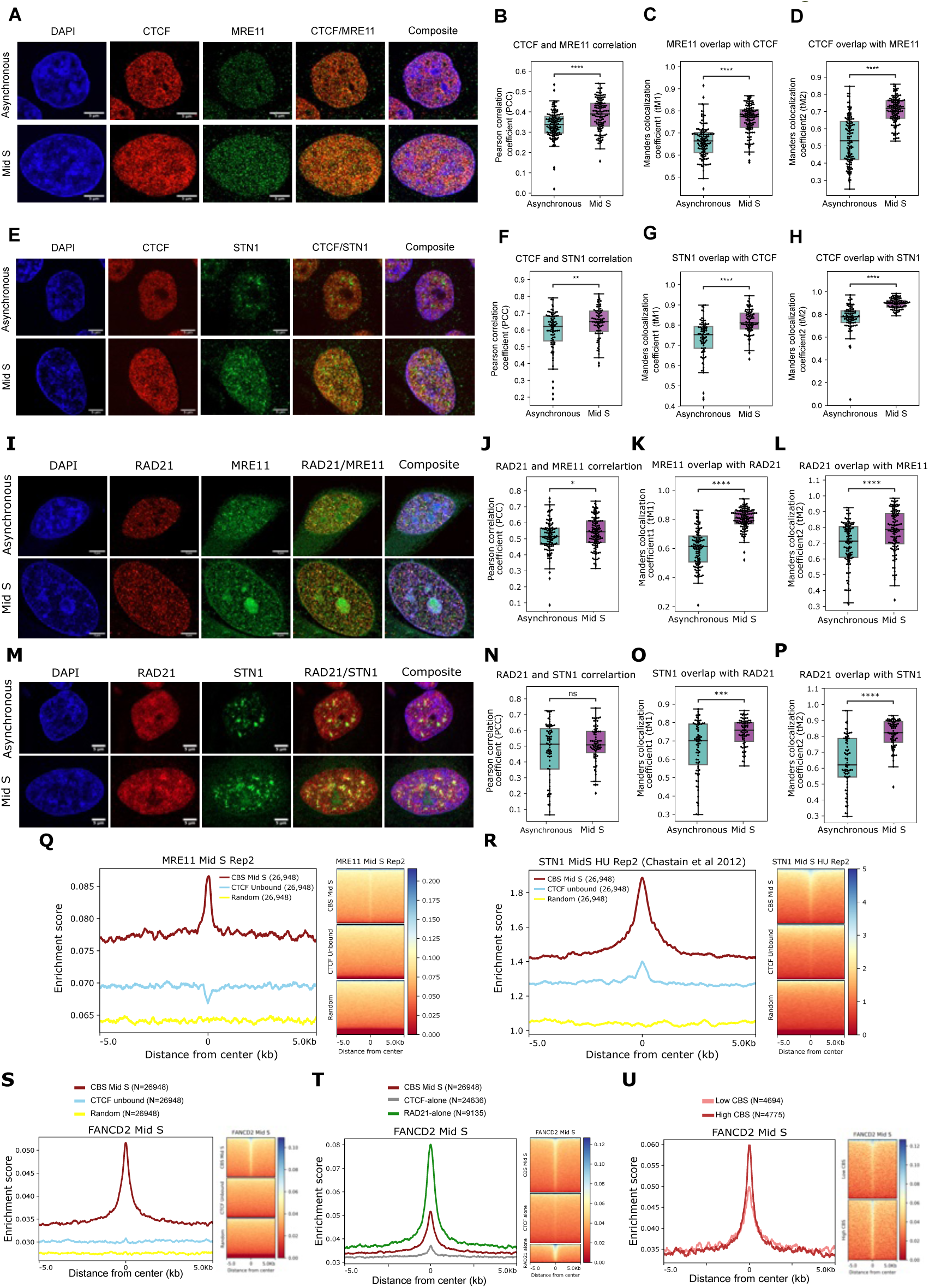
Enrichment of replication stress-associated proteins at CTCF/cohesin-binding sites. **(A)** Representative images of asynchronous and Mid S HeLa cells co-immunostained with CTCF (red, Alexa Fluor-568) and MRE11 (green, Alexa Fluor-488). DNA is counter-stained with DAPI (blue). Images were acquired with ZEISS LSM 980 with Airyscan 2 microscope in super-resolution (SR) mode. Scale bar, 5µm. No. of cells n=135, and No. of biological replicates N=3. Quantification of nuclear colocalization between CTCF and MRE11 by **(B)** Pearson’s Correlation Coefficient (PCC: P=2.21x10-07) and Mander’s overlap coefficients **(C)** tM1 for MRE11 i.e., the fraction of MRE11 overlapping with CTCF (P=1.05x10-22) and **(D)** tM2 for CTCF i.e., the fraction of CTCF overlapping with MRE11 (P=1.44x10-24). **(E)** CTCF co-stained with STN1 in asynchronous and S-phase synchronised cells. No. of cells n=88, and No. of biological replicates N=3. Colocalization quantified by **(F)** PCC (P= 6.99x10-03) and Mander’s **(G)** tM1 (P=2.35x10-10) and **(H)** tM2 (P=4.31x10-19). **(I)** Representative images of asynchronous and Mid S HeLa cells co-immunostained with RAD21 and MRE11 from three biologically independent experiments. No. of asynchronous cells=118, No. of Mid S cells=129. Quantification of nuclear colocalization between RAD21 and MRE11 by **(J)** PCC (P=3.24x10^-02^) and Mander’s overlap coefficients **(K)** tM1 for RAD21 (P=1.28x10^-29^) and **(L)** tM2 for MRE11 (P=1.12x10^-5^). **(M)** Representative images of RAD21 co-stained with STN1 in asynchronous and S-phase synchronised cells. No. of cells n=79, No. of biological replicates N=2. Colocalization quantified by **(N)** PCC (P=4.1x10^-01^) and Mander’s **(O)** tM1 (P=6.02x10^-4^) and **(P)** tM2 (P=1.44x10^-12^). Colocalization was quantified using the JACoP plugin in Fiji. DAPI staining was used to draw ROI to get only the nuclear signals. The threshold signal was determined with Costes’ automatic threshold method and manual adjustment. Centerline: median; Box limits: 1st and 3rd quartile; whiskers: 1.5xIQR. p-values were determined by a two-sided non-parametric Mann-Whitney test. p-value annotation legend: ns: 5.00x10^-02^ < p <= 1, *: 1.00x10^-02^ < p <= 5.00x10^-02^, **: 1.00x10^-03^ < p <= 1.00x10^-02^, ***: 1.00x10^-04^ < p <= 1.00x10^-03^, ****: p <= 1x10^-04^. **(Q)** Second biological replicate of MRE11 ChIP-seq signal in Mid S HeLa cells plotted across a +/-5kb window centred at Mid S CBSs (maroon), CTCF unbound sites (sky blue) and random genomic regions (yellow). 26,948 sites were in each category. **(R)** ChIP-seq profile of STN1 (second biological replicate) in Mid S upon HU treatment from Chastain et al. [32] plotted around CBSs and control sites (+/-5kb). **S-U:** ChIP-Seq signal of FANCD2 in Mid S HeLa cells plotted at **(S)** CBS and control sites, **(T)** CBS, CTCF-alone and RAD21 alone sites and **(U)** low and high-binding CBS.

**Fig S3.**
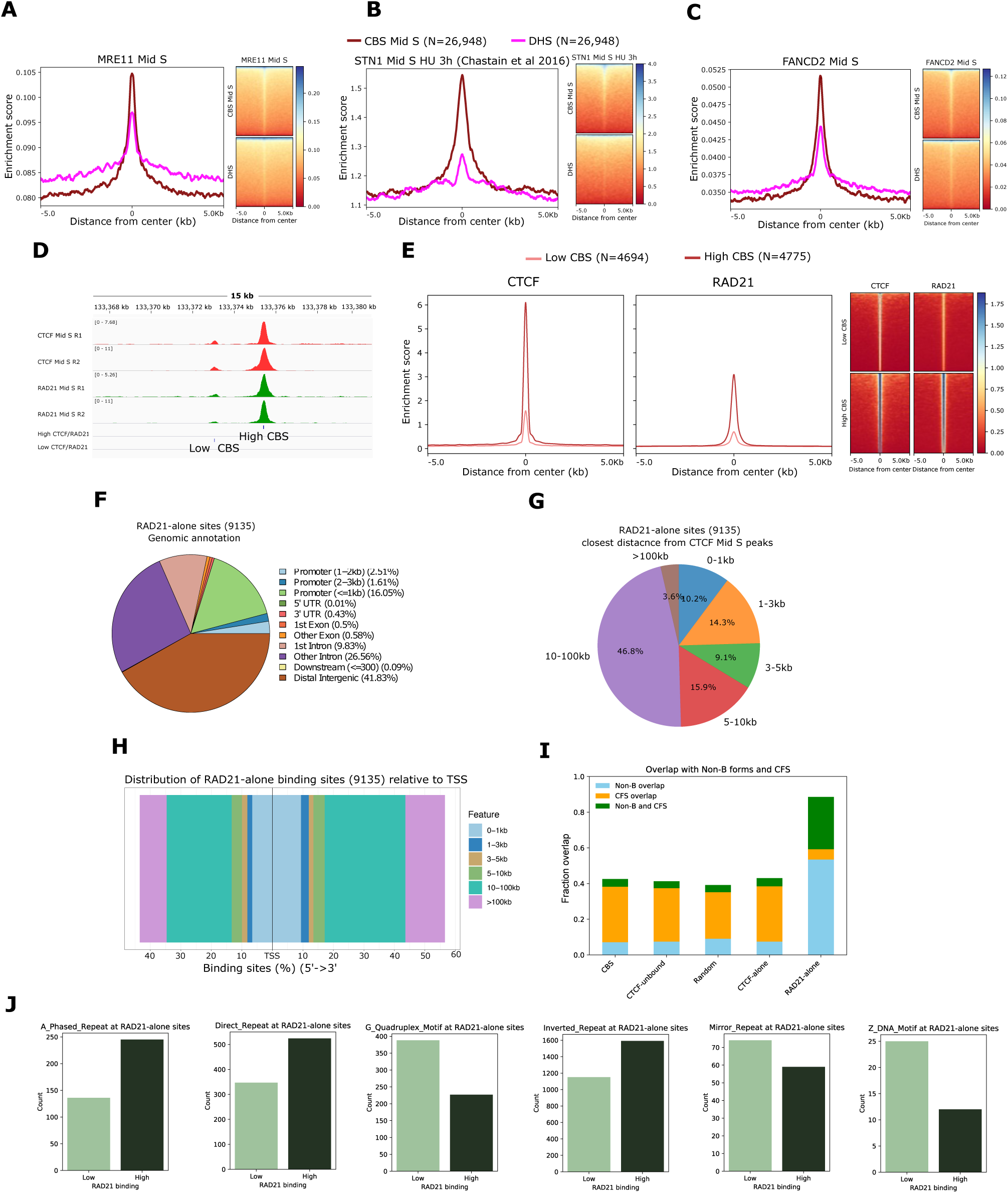
ChIP-Seq enrichment and RAD21-alone sites. A-C: Enrichment of **(A)** MRE11 **(B)** STN1 (+HU) and **(C)** FANCD2 plotted at a +/-5kb window of CBSs (maroon) and DHSs without overlapping CTCF/RAD21 peaks (magenta). **(D)** Genome browser snapshot of two CBSs showing low- or high-binding of CTCF and RAD21. **(E)** ChIP-seq profile and heatmap of CTCF and RAD21 at low- and high-binding sites. **(F)** Genomic annotation of RAD21-alone sites (9135). **(G)** Distribution of closest distance of RAD21-alone sites relative to CTCF Mid S peaks (87734). **(H)** Distribution of RAD21 alone-sites from transcription start sites (TSS). **(I)** Fractions of genomic sites overlapping with non-B DNA structures and human chromosomal fragile sites (CFS). 26,948 sites are present in CBS, CTCF-unbound and random regions. CTCF-alone and RAD21-alone sites consist of 24,636 and 9,135 sites, respectively. **(J)** Presence of non-B DNA repeats (inverted repeats, G-quadruplex motifs, direct repeats, short-tandem repeats, A-phased repeats, mirror repeats and z-DNA motifs) at low binding and high binding RAD21-alone sites. 3045 sites are present in both the sets.

**Fig S4.**
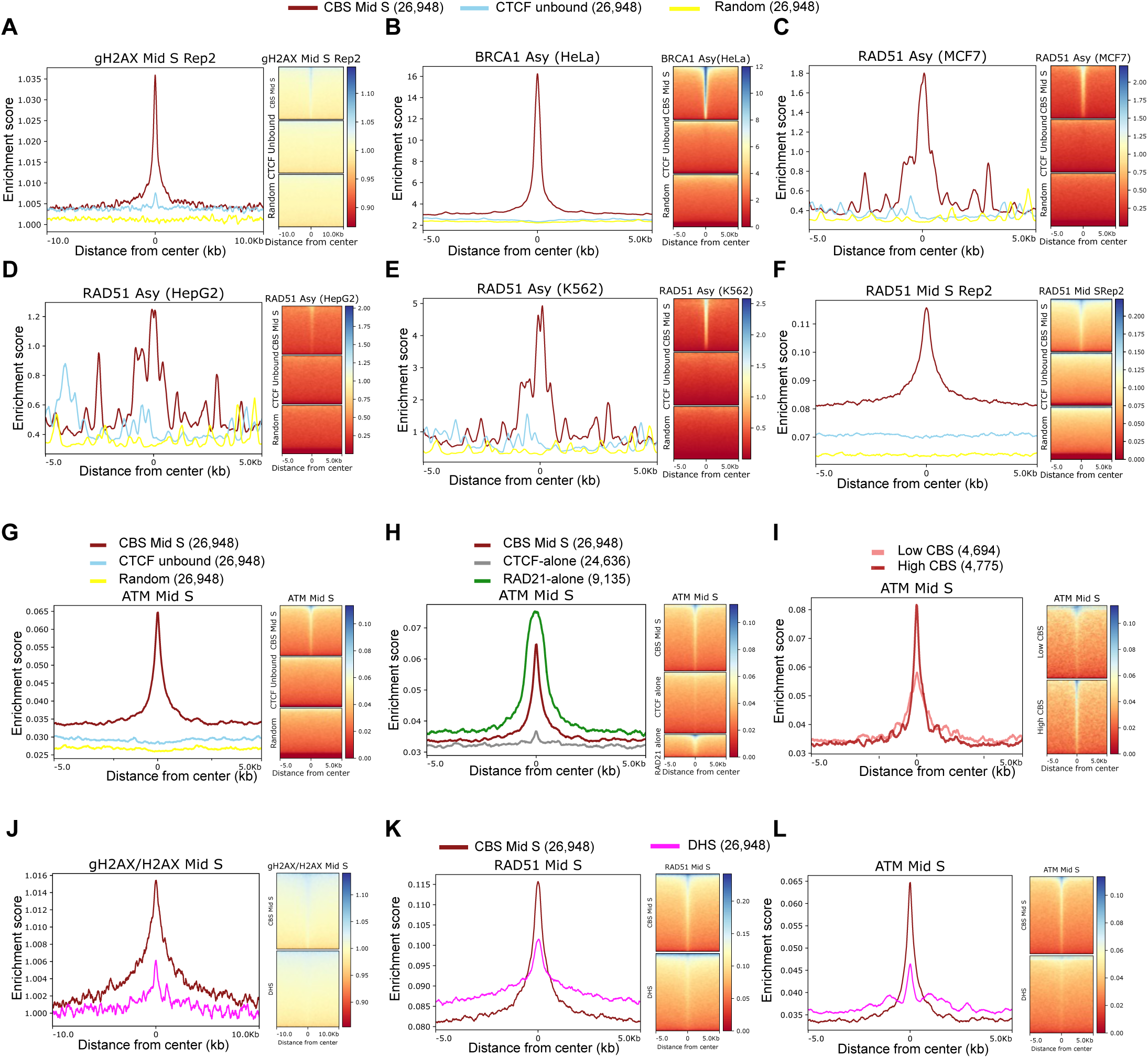
Enrichment of γH2AX, BRCA1 and RAD51 ChIP-seq signals at CBSs. **(A)** γH2AX ChIP-seq signal in Mid S HeLa cells from second biological replicate plotted at CBS Mid S, CTCF unbound sites and random sites. **(B)** BRCA1 ChIP-seq signal from the asynchronous population of HeLa cells downloaded from ENCODE plotted at CBS and control sites. RAD51 ChIP-seq from ENCODE **(C)** MCF7 **(D)** HepG2 and **(E)** K562cells plotted at CBSs and control sites in asynchronous. **(F**) Second replicate of RAD51 ChIP-seq signal from Mid S synchronised HeLa cells at CBS Mid S and control sites. **G-I:** ChIP-Seq signal of FANCD2 in Mid S HeLa cells plotted at **(G)** CBS and control sites, **(H)** CBS, CTCF-alone and RAD21 alone sites and **(I)** low and high-binding CBS. **J-L:** Enrichment of **(J)** gH2AX (Normalised with H2AX) **(K)** RAD51 and **(L)** ATM plotted at a +/-5kb window of CBSs (maroon) and DHSs without overlapping CTCF/RAD21 peaks (magenta).

**Fig S5.**
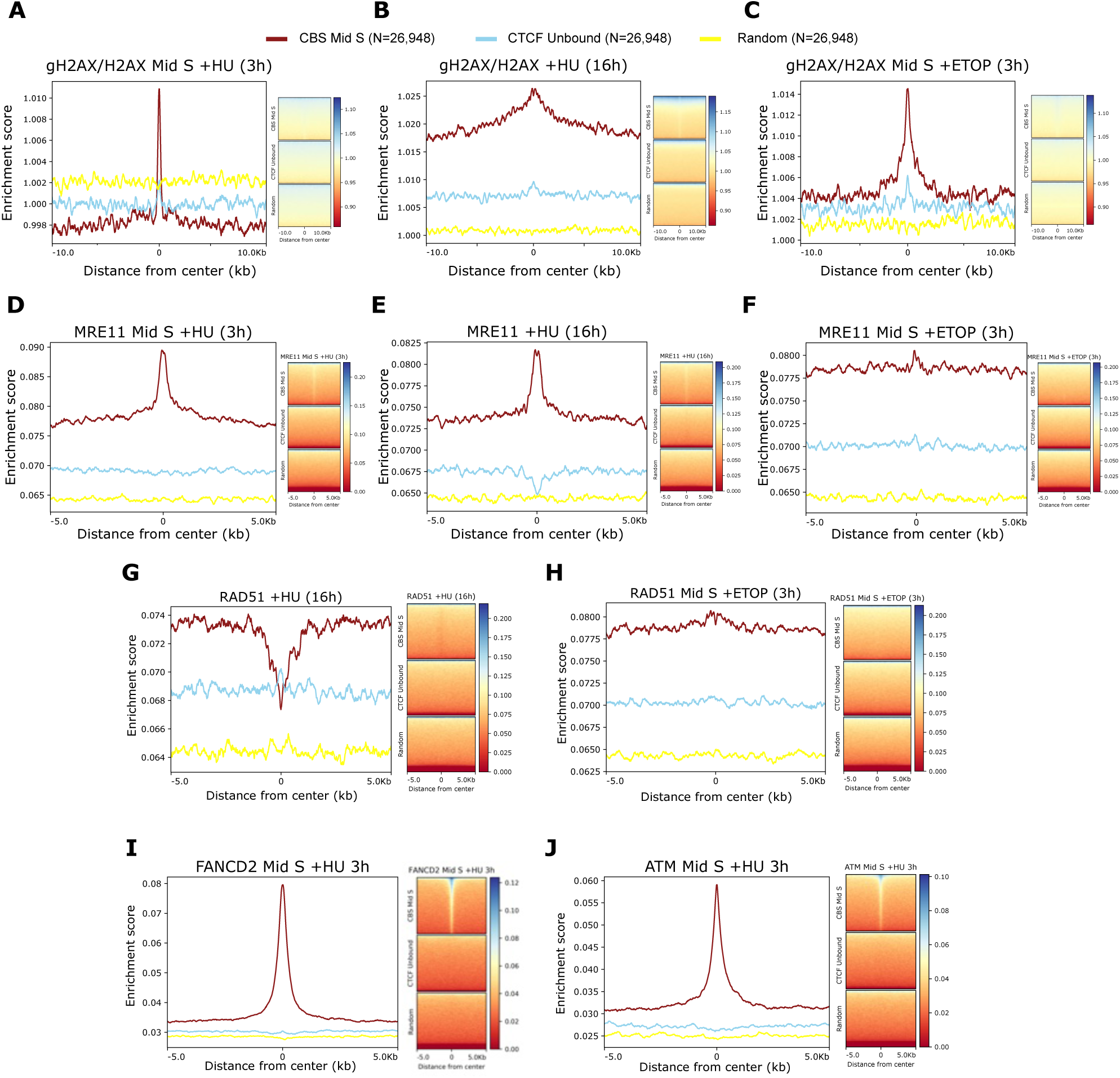
ChIP-seq signal of DDR proteins in exogenous replication stress conditions. A-C: ChIP-seq profile and heatmap of γH2AX (normalised with H2AX) from **(A)** Mid S HeLa cells treated with 2 mM HU for three hours **(B)** asynchronous HeLa cells treated with 2 mM HU for 16 hours and **(C)** Mid S HeLa cells treated with 25µM Etoposide for 3 hours plotted at CBSs (26,948 sites), CTCF unbound (26,948 sites) and random regions (26,948 sites). **D-F:** ChIP-seq profile and heatmap of MRE11 from **(D)** Mid S HeLa cells treated with 2 mM HU for three hours **(E)** HeLa cells treated with 2 mM HU for 16 hours and **(F)** Mid S HeLa cells treated with 25 µM Etoposide for 3 hours plotted at CBSs, CTCF unbound and random sites. **G-H:** ChIP-seq profile and heatmap of RAD51 from **(G)** HeLa cells treated with 2 mM HU for 16 hours and **(H)** Mid S HeLa cell treated with 25 µM Etoposide for 3 hours at CBSs and control sites. **I-J:** ChIP-Seq profile of **(I)** FANCD2 and **(J)** ATM from Mid S HeLa cells treated with 2 mM HU for three hours.

**Fig S6.**
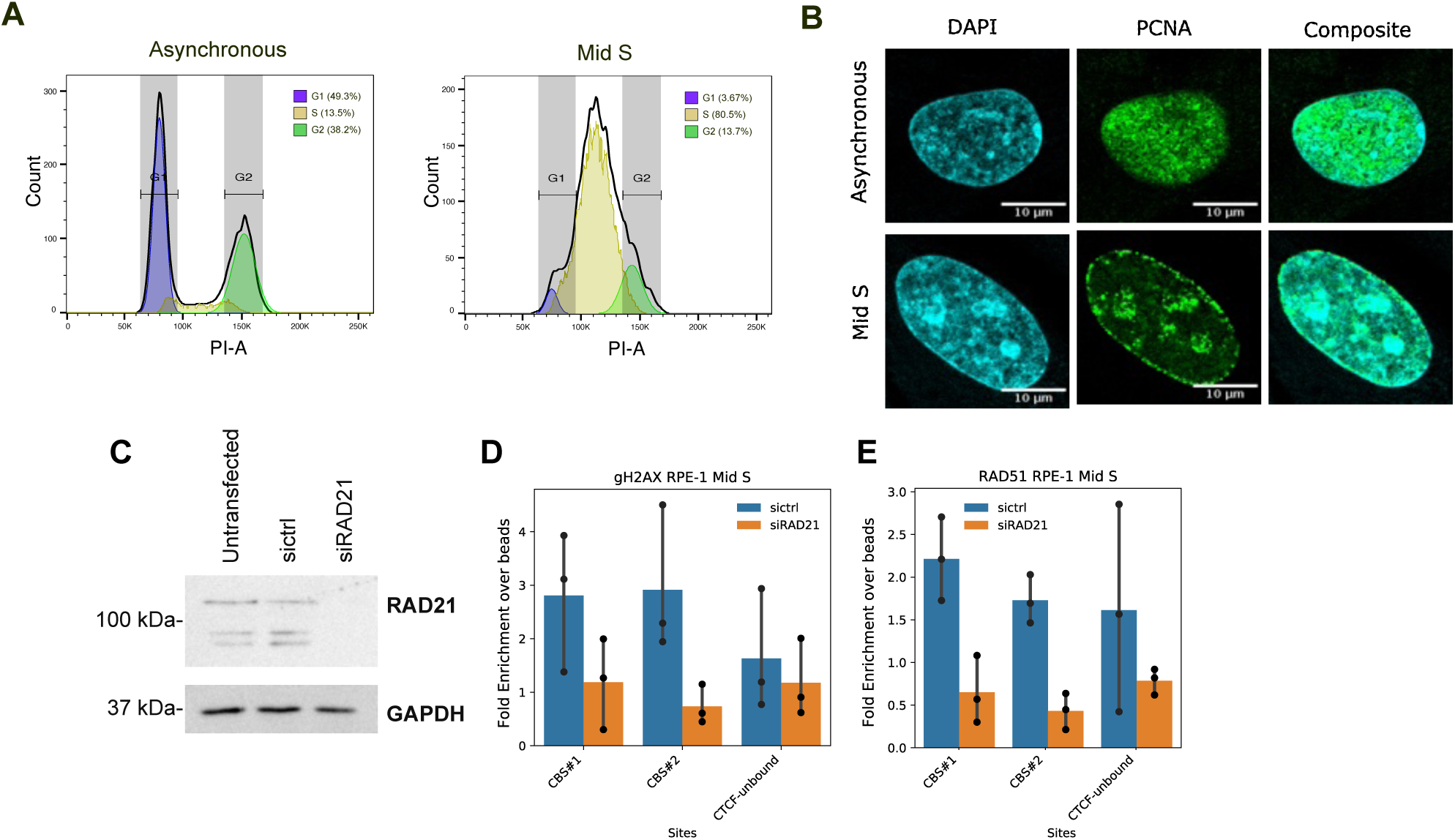
ChIP in hTERT RPE-1 cells in Mid S phase. **(A)** Flow Cytometry analysis of PI stained asynchronous and Mid S synchronised hTERT RPE-1 cells.The percentage of cells in each phase of the cell cycle is mentioned in brackets. **(B)** Representative immunostaining images of PCNA (green) in hTERT RPE-1 cells. The nucleus is counterstained with DAPI (blue). Mid S cells are showing punctate of PCNA at nuclear and nucleolar boundaries. The scale bar is 10µM. **(C)** Western blotting of RAD21 in untransfected and siRNA transfected hTERT RPE-1 cells synchronized in Mid S phase confirming RAD21 knockdown. GAPDH is used as a loading control. **D,E:** ChIp-qPCR of **(D)** gH2AX and **(E)** RAD51 in Mid S synchronized hTERT RPE-1 cells with control siRNA or siRAD21 transfection at two CBSs and a CTCF-unbound site (from 3 replicates). The y-axis (fold enrichment over beads) indicates the % input in immunoprecipitation (IP) divided by that of beads.

**Fig S7.**
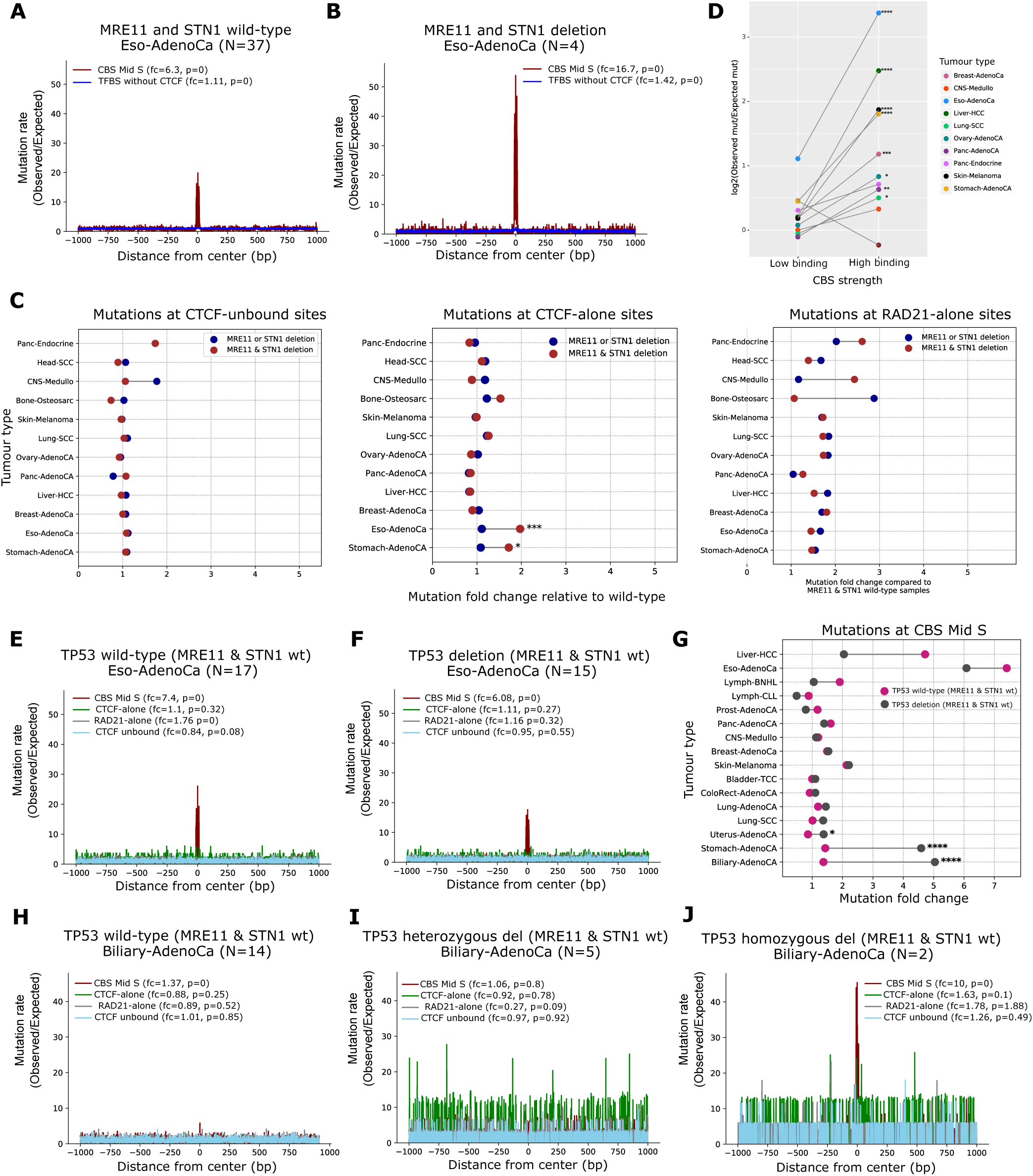
Somatic mutation analysis. A,B: Mutation rate plotted at +/-1 kb region of CBS Mid S and Transcription factor binding sites (TFBS excluding CTCF and RAD21) in Esophageal adenocarcinoma samples with **(A)** both MRE11 and STN1 wild-type (No. of samples N=37) **(B)** both MRE11 and STN1 deletion sample (N=4). P-value is computed by G-test. **(C)** Mutation ratio between MRE11 or/and STN1 deletion to MRE11 and STN1 wild-type samples at CTCF unbound sites, CTCF-alone sites and RAD21-alone sites across tumour types. **(D)** Mutation rate at low and high binding CTCF/cohesin sites (+/-20 bp) computed in various tumour types based on MRE11-STN1 deletion status. The p-value was calculated using Fisher’s exact test. **E, F:** Mutation rate at CBSs and control sites in **(E)** wild-type TP53 (N=17) and **(F)** TP53 deletion samples (N=15) from Esophageal adenocarcinoma with no alterations in MRE11 and STN1 (i.e., MRE11 and STN1 wild-type). **(G)** Mutations fold change at CBSs in TP53 wild-type and deletion samples from different tumour types. **H, I, J:** Mutation rate plotted at CBS Mid S and control sites in Biliary adenocarcinoma samples (with background of wild-type MRE11 and STN1) with **(H)** TP53 wild-type (N=14) **(I)** TP53 deletion in heterozygous condition (N=5) **(J)** TP53 deletion in homozygous condition (N=2).

## Notes

### Competing Interest Statement

The authors have declared no competing interest.

### Summary of Updates

Additional experiments have been included to support our conclusions further. a) ChIP-seq of additional replication-stress marker proteins have been included (Fig. S2, S4 and S5). b) DNA damage response at CTCF/Cohesin-binding sites in normal cell lines have been studied and compared with the cancerous cell lines (Fig. 4 and Fig S6).

